# Bumetanide Strengthens Residual Brain–Bladder Communication After Spinal Cord Injury

**DOI:** 10.64898/2026.08.28.747863

**Authors:** Qiang Li, Wei Li, Junkui Shang, Alfredo Sandoval, Tiffany Dunn, Hee Young Kim, Kathleen L. Vincent, Bo Chen

## Abstract

Neurogenic bladder is one of the most disabling consequences of spinal cord injury (SCI), yet it remains unclear whether residual brain–bladder communication persists after injury and can be therapeutically strengthened. Here, we integrated analysis of a clinical SCI cohort with studies in a mouse model that recapitulates key features of human neurogenic bladder. SCI markedly impaired both descending and ascending limbs of the spinobulbospinal micturition reflex, reducing bladder responses evoked by stimulation of the pontine micturition center (PMC) and bladder filling-induced activation of the periaqueductal gray. Despite this marked functional impairment, pseudorabies virus tracing, optogenetics, and machine learning-assisted three-dimensional imaging revealed persistent bladder-connected neurons within the spared T8–T9 interlesion region. Neurons within this region remained responsive to descending PMC input, identifying the interlesion network as a candidate substrate for residual brain–bladder communication. Early continuous intrathecal bumetanide improved urinary storage and emptying, enhanced descending PMC-to-bladder and ascending bladder-to-brain signaling, and increased the functional engagement of bladder-connected T8–T9 neurons without increasing urine production. Bumetanide also increased trans-synaptic labeling between the bladder and PMC, consistent with enhanced polysynaptic brain–bladder connectivity. Together, these findings indicate that brain–bladder communication is not completely lost after SCI and may be supported by a functionally compromised residual spinal network that remains amenable to therapeutic reinforcement. Targeting spared autonomic circuitry may therefore provide a strategy for improving bladder function after SCI.

**Highlights:**

- SCI disrupts bidirectional communication within the brain–bladder circuit.
- Spared T8–T9 interlesion neurons remain engaged in residual brain–bladder signaling.
- Intrathecal bumetanide strengthens brain–bladder signaling and improves urinary function.

**Graphic abstract:** 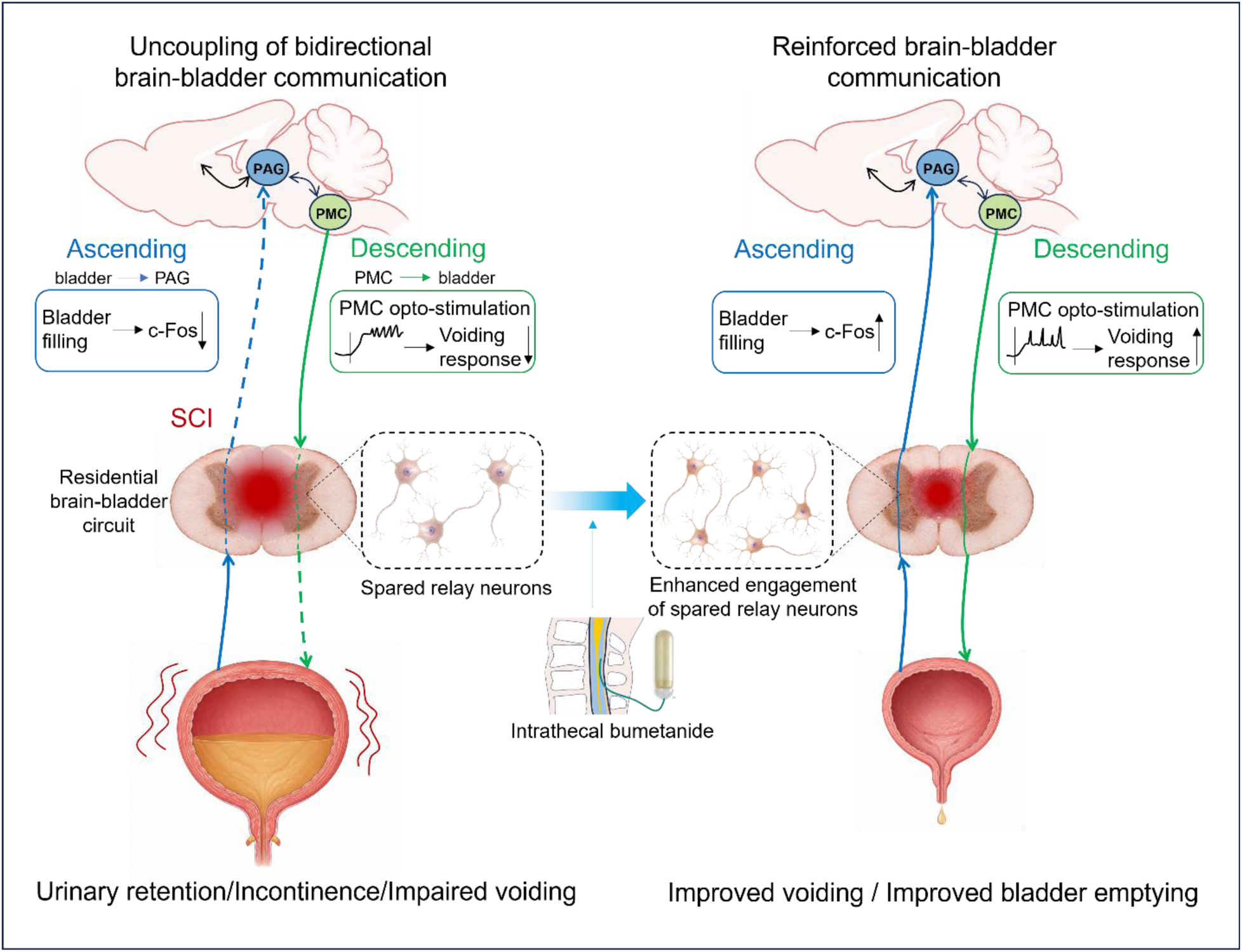

## INTRODUCTION

Spinal cord injury (SCI)–associated neurogenic bladder affects more than 80% of patients and remains a major cause of morbidity, recurrent hospitalization, loss of independence, and reduced quality of life^1–3^. Common complications include urinary tract infection, urinary stones, renal impairment, incontinence, and high-pressure bladder storage^2^. Current management, including clean intermittent catheterization, antimuscarinics, botulinum toxin, and surgical or device-based interventions, can reduce complications and improve bladder storage or emptying. However, these treatments do not restore coordinated supraspinal control of the storage-to-voiding transition. Consistent with this unmet need, clinical and urodynamic assessment of our SCI cohort revealed broad impairment across both storage and voiding domains.

Efficient micturition requires coordinated bidirectional communication between the bladder and brain through the spinobulbospinal reflex, which links bladder afferents, sacral spinal circuits, the periaqueductal gray (PAG), and the pontine micturition center (PMC)^4–6^. During bladder filling, ascending sensory signals activate PAG networks that integrate bladder fullness and engage descending PMC output to coordinate parasympathetic and sphincter circuits for efficient voiding^4,7^. Following SCI, this bidirectional communication is profoundly disrupted. Loss of supraspinal regulation together with maladaptive spinal reflexes leads to detrusor overactivity, detrusor–sphincter dyssynergia, elevated bladder pressure, and inefficient bladder emptying^4,8^. However, whether brain–bladder communication is completely abolished or whether residual supraspinal communication persists after SCI remains unknown.

Studies of locomotor recovery have shown that spared propriospinal circuits can relay descending commands across incomplete spinal lesions^9,10^. These findings raise the possibility that analogous intraspinal circuits may preserve residual communication between supraspinal centers and the lower urinary tract. Whether such relays persist after SCI, participate in bladder control, and remain amenable to therapeutic reinforcement has not been established.

We previously showed that a staggered double lateral hemisection SCI model induces prolonged neuronal swelling and loss of excitatory neurons within the spared interlesion region, contributing to functional dormancy of descending pathways. Intrathecal bumetanide reduced neuronal swelling, preserved interlesion neurons, and improved locomotor recovery^11^. Bumetanide is a potent loop diuretic that penetrates the central nervous system poorly after systemic administration, intrathecal delivery may therefore increase spinal exposure while limiting renal effects^12–14^. We hypothesized that protecting vulnerable neurons within the interlesion region would preserve relay transmission and strengthen residual communication between the brain and bladder.

Here, we combined clinical phenotyping with a staggered mouse model of SCI to determine whether residual brain–bladder communication persists after injury and can be therapeutically strengthened. Using optogenetic circuit interrogation, transsynaptic tracing from the bladder, three-dimensional circuit mapping, and pharmacological intervention, we show that SCI disrupts both the descending and ascending limbs of the spinobulbospinal micturition reflex. We further identify a spared thoracic interlesion network that retains residual connectivity with supraspinal bladder-control circuits and demonstrate that early intrathecal bumetanide strengthens functional brain–bladder communication and enhances engagement of the spared interlesion network.

## RESULTS

### Spinal cord injury produces a severe and persistent neurogenic bladder phenotype in humans and mice

To define the clinically relevant features that a preclinical model should recapitulate, we characterized urinary dysfunction in a cohort of patients with chronic SCI (Fig. 1A; Table 1). The cohort included 17 patients evaluated a mean of 15 ± 12 years after injury, encompassing both complete and incomplete SCI with cervical, thoracic, and lumbar lesions. All patients required clean intermittent catheterization (CIC) for bladder management. Despite standard care, patients continued to exhibit substantial neurogenic bladder symptoms. Neurogenic Bladder Symptom Score (NBSS) profiling demonstrated persistent storage/voiding dysfunction together with urinary incontinence (Fig. 1B; Fig. S1A). Consistently, American Urological Association (AUA) Symptom Score remained elevated (14.6 ± 6.4; Fig. 1C), indicating persistent lower urinary tract symptoms. Objective clinical measures further demonstrated impaired bladder emptying and urinary incontinence, as reflected by increased post-void residual (PVR) volumes and frequent urinary leakage episodes (Fig. 1D). A representative urodynamic tracing showed elevated bladder pressure during filling, urinary leakage preceding coordinated voiding, and impaired bladder emptying (Fig. S1B). Neurogenic bladder symptoms also imposed a substantial burden on patient quality of life (Fig. S1C).

**Figure 1.**
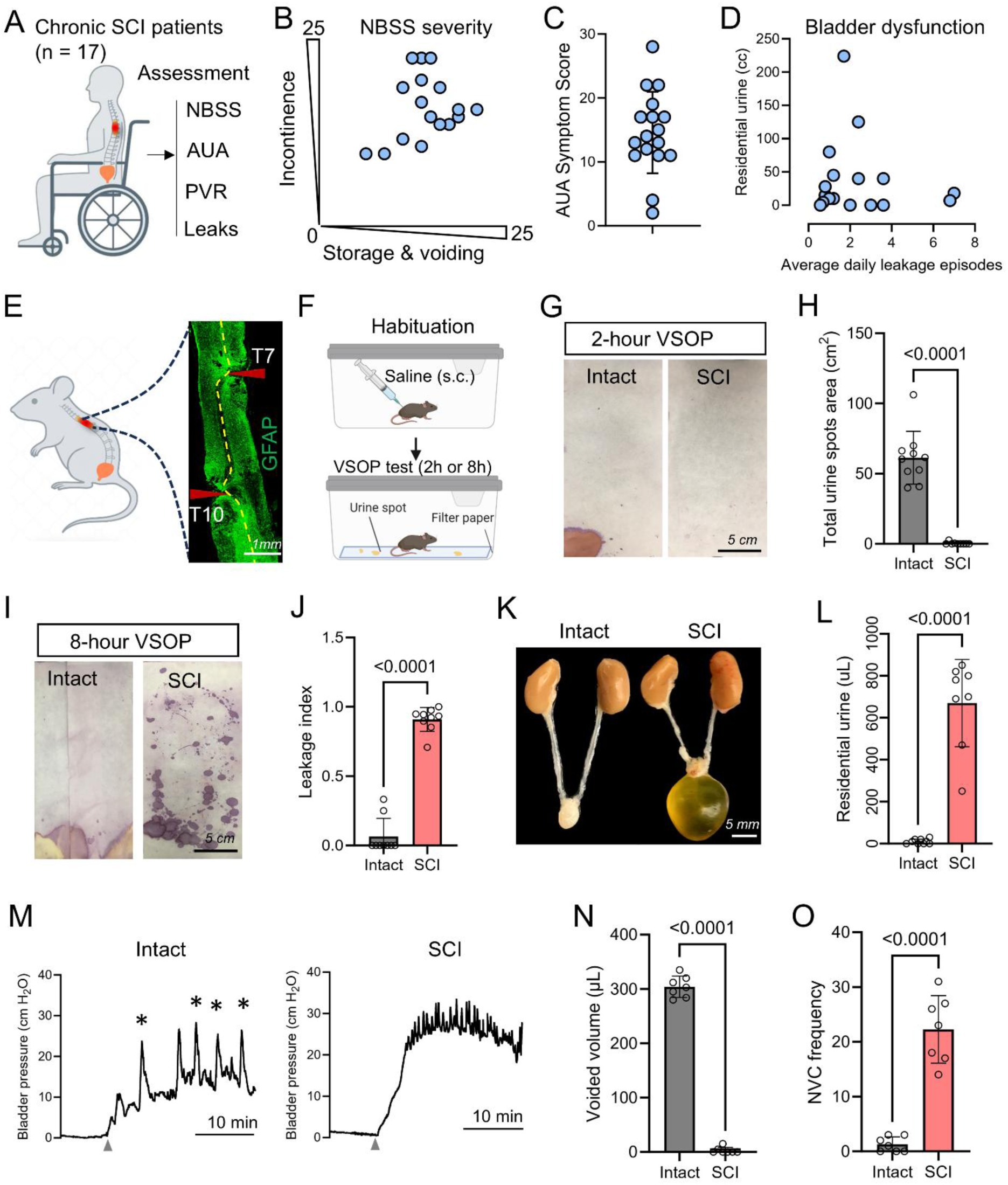
Clinical burden and mouse modeling of neurogenic bladder after SCI. (A) Schematic of the chronic SCI cohort (n = 17) and clinical assessments, including the Neurogenic Bladder Symptom Score (NBSS), American Urological Association Symptom Score (AUA), post-void residual urine (PVR), and urinary leakage frequency. (B) Individual NBSS profiles showing storage and voiding symptom severity versus incontinence severity in patients with chronic SCI. (C) Individual AUA Symptom Scores in the SCI cohort. (D) Individual patient distribution of residual urine volume and average daily urinary leakage episodes. (E) Staggered SCI mouse model with hemisections at T7 and T10; representative GFAP staining shows the two lesion sites and preserved interlesion spinal segment. (F) Experimental workflow for the voiding spot on paper (VSOP) assay following habituation and subcutaneous saline administration. (G, H) Representative 2-h VSOP images and quantification of total urine spot area in intact and SCI mice. N = 10, Unpaired t test with Welch’s correction. (I, J) Representative 8-h VSOP images and quantification of leakage index. N = 8-9, Unpaired t test with Welch’s correction. (K, L) Representative gross urinary tract morphology and quantification of residual urine volume in intact and SCI mice. N = 7-8, Unpaired t test with Welch’s correction. (M) Representative conscious cystometric recordings from intact and SCI mice. Arrowheads indicate the onset of bladder filling, and asterisks indicate voiding contractions. (N, O) Quantification of voided volume and non-voiding contraction (NVC) frequency during conscious cystometry. N = 7, Unpaired t test with Welch’s correction. Data are presented as mean ± SD. Each point represents an individual participant or mouse, as appropriate. Scale bars, 1 mm (E), 5 cm (G, I), and 5 mm (K).

**Table 1.**
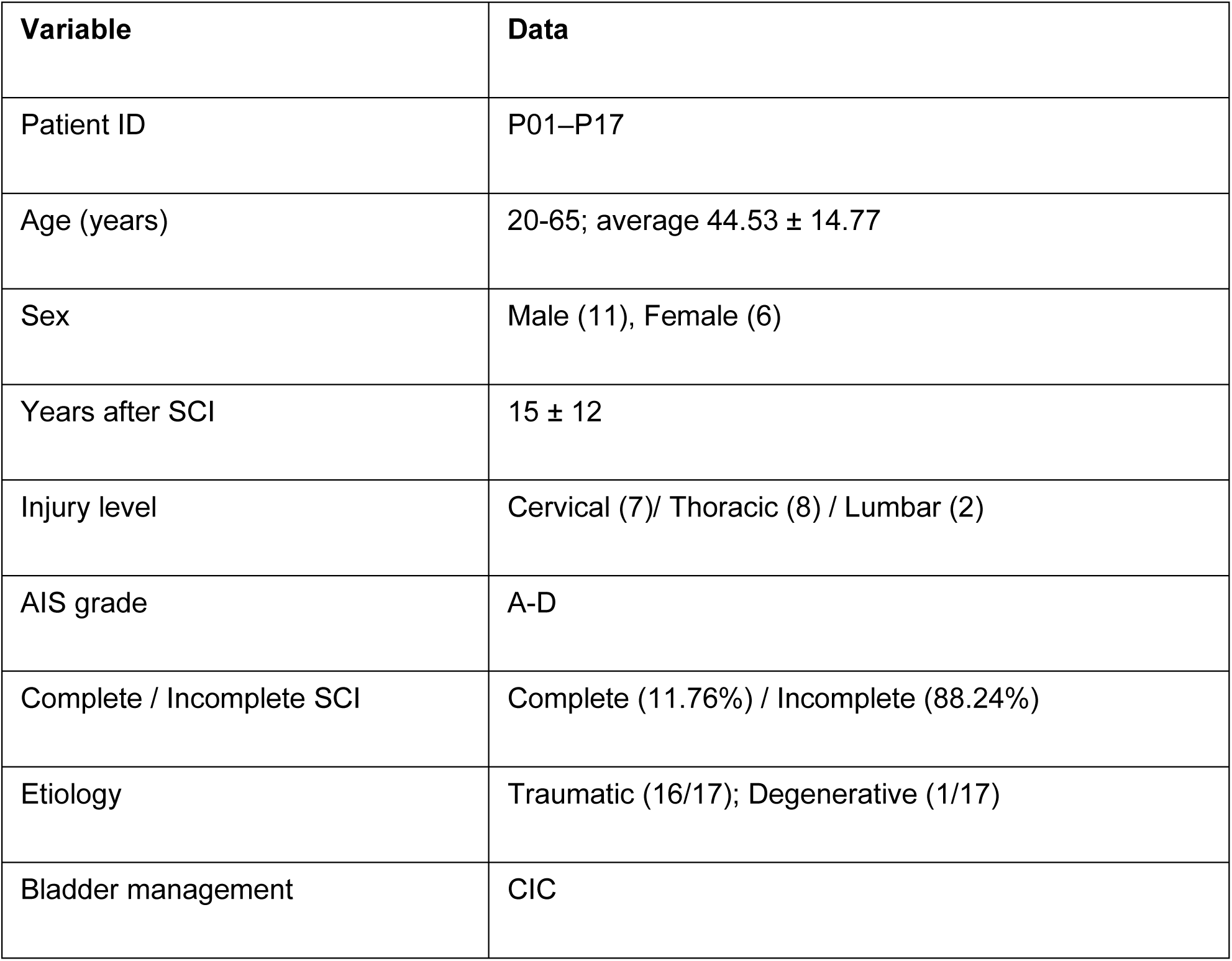
Summary of patients in our cohort.

To establish an experimental model that recapitulates the clinical features of neurogenic bladder after SCI, we performed staggered lateral hemisections at T7 and T10 (Fig. 1E), disrupting descending supraspinal pathways while preserving a narrow bridge of spared spinal tissue between the lesions^9,11,15^. SCI mice developed profound defects in coordinated bladder storage and voiding. In the voiding spot on paper (VSOP) assay, intact mice generated large, spatially restricted urine spots within 2 h, whereas SCI mice exhibited complete urinary retention with an absence of detectable voiding events (Fig. 1F–H). By 8 h after injury, SCI mice developed fragmented voiding characterized by numerous small, scattered urine spots, consistent with urinary leakage and loss of coordinated voiding (Fig. 1I–J; Fig. S2A–B). These abnormalities were accompanied by marked bladder distension and substantially increased residual urine volumes (Fig. 1K–L).

Conscious cystometry further confirmed severe bladder dysfunction following SCI. Representative cystometric recordings demonstrated a transition from coordinated filling–voiding cycles in intact mice to sustained bladder overactivity after SCI, characterized by markedly reduced voided volume and frequent non-voiding contractions (Fig. 1M–O). Quantitative analysis further demonstrated reduced confirmed voiding frequency, increased residual bladder volume, increased functional bladder capacity, and elevated mean filling pressure (Fig. S2E–H). Together, these findings indicate impaired bladder emptying, abnormal storage-phase activity, and inefficient voiding following SCI, closely paralleling the major clinical features observed in patients with chronic SCI.

### Staggered SCI disrupts both arms of the spinobulbospinal micturition reflex

Because coordinated micturition depends on bidirectional signaling between the PMC and sacral spinal circuits^4,5^ (Fig. 2A), we asked whether staggered SCI disrupts both arms of the spinobulbospinal micturition reflex.

**Figure 2.**
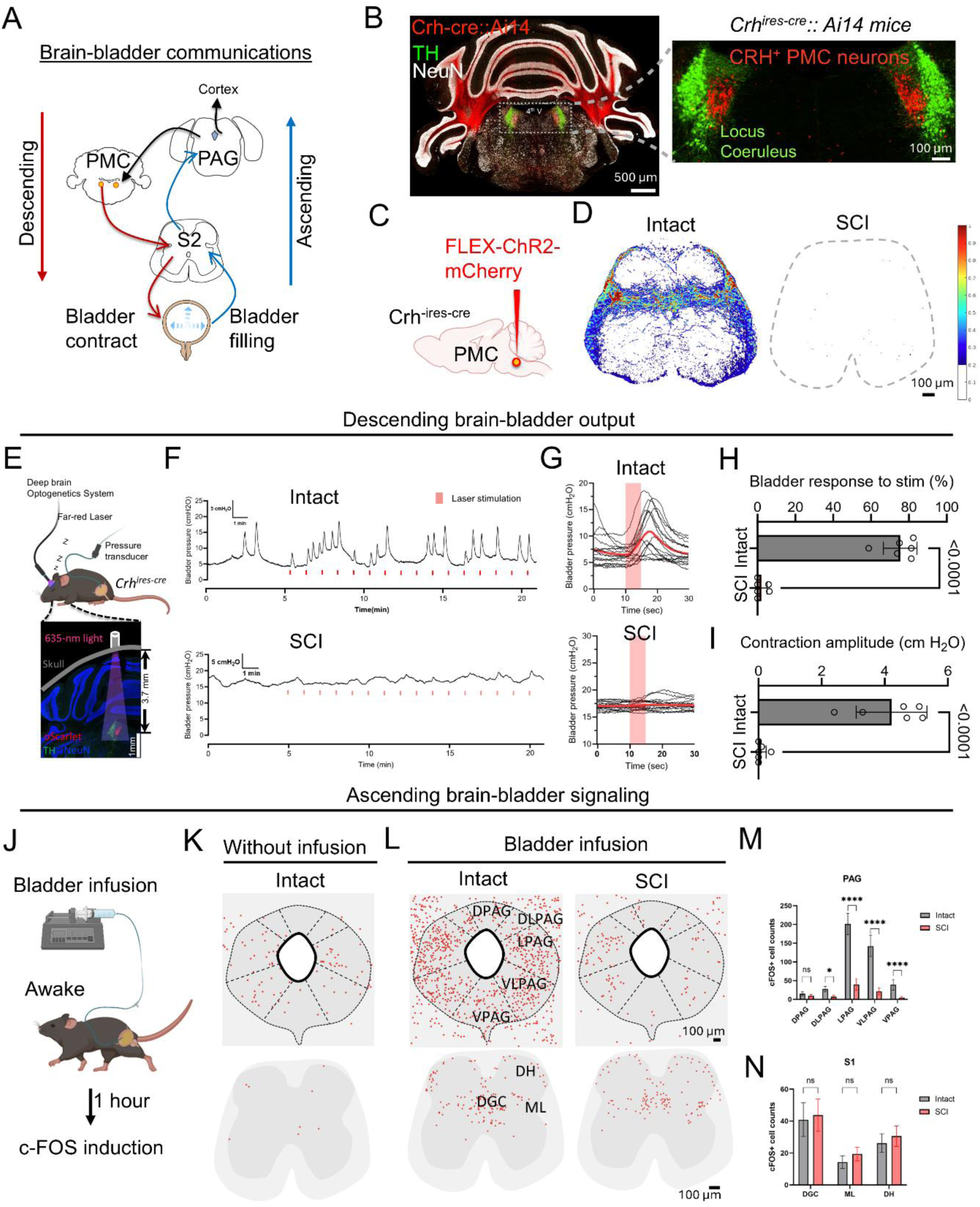
Bidirectional interrogation of brain–bladder communication after SCI. (A) Schematic of descending PMC control of bladder contraction and ascending bladder-filling signals to PAG/PMC. (B) Representative sections from *Crh*^ires-Cre^::Ai14 mice showing CRH^+^ PMC neurons and their localization relative to the locus coeruleus. (C) Viral strategy for ChR2-mediated mapping of CRH^+^ PMC neurons. (D) Representative S2 spinal cord projection heatmaps showing loss of descending PMC input after SCI. (E) Experimental setup for non-invasive optogenetic stimulation of PMC CRH^+^ neurons with simultaneous bladder pressure recording. (F, G) Representative and aligned bladder pressure traces during laser stimulation in intact and SCI mice. (H, I) Quantification of stimulation-evoked bladder response rate and contraction amplitude. N = 6, Unpaired t test with Welch’s correction. (J) Experimental design for awake bladder infusion and c-Fos induction. (K, L) Representative c-Fos maps without infusion and after bladder infusion in PAG and PMC regions of intact and SCI mice. (M, N) Quantification of c-Fos^+^ cells in PAG and PMC subregions. N = 6, Two-way ANOVA followed by Bonferroni’s multiple comparisons test. Data are shown as mean ± SD. Scale bars as indicated.

To characterize the descending arm anatomically, we first localized PMC Crh^+^ neurons in coronal brainstem sections from Crh^Ires-Cre^; Ai14 reporter mice. Consistent with previous studies^6,16^, tdTomato-labeled Crh^+^ neurons were clustered medial and adjacent to the locus coeruleus (Fig. 2B). We next injected AAV2/8-Flex-ChR2-mCherry into the PMC of Crh^Ires-Cre^ mice to trace descending projections from PMC Crh^+^ neurons (Fig. 2C; Fig. S3A). In intact mice, Crh^+^ axons robustly innervated the sacral parasympathetic region, whereas staggered SCI abolished detectable descending projections at sacral levels (Fig. 2D; Fig. S3B), indicating anatomical disconnection of the PMC–sacral bladder pathway.

We next tested whether this pathway remained functionally coupled to bladder output after SCI. To this end, we combined non-invasive optogenetic stimulation^17^ of PMC Crh^+^ neurons with cystometry (Fig. 2E; Fig. S3E). Anesthetized Crh^Ires-Cre^ mice received AAV-nEF-Con-Foff2.0-ChRmine-oScarlet or control virus targeting PMC Crh^+^ neurons, and intravesical pressure was recorded during repeated far red-light stimulation (Fig. 2F). In intact ChRmine mice, each train elicited a rapid, time-locked bladder contraction superimposed on the filling rhythm (Fig. 2G). The same stimulus failed to elicit bladder contractions in SCI mice or in intact control-virus mice (Fig. 2G; Fig. S3C-D). Light trains produced bladder contraction on ∼76% of trials in intact ChRmine mice versus <5% in controls and SCI (Fig. 2H), and the evoked contraction amplitudes were markedly reduced after SCI (Fig. 2I). These findings demonstrate that Crh^+^ PMC neurons are sufficient to drive bladder contraction in intact mice and that staggered SCI abolishes functional descending PMC–bladder communication.

We next examined the ascending arm of the reflex by assessing bladder filling–evoked activation in the periaqueductal gray (PAG) and PMC, as the PAG is a major supraspinal relay that receives bladder afferent input and transmits it to the brain and PMC to coordinate micturition^7,18^. Intact and 8-week post-SCI mice underwent continuous-infusion conscious cystometry, followed by c-Fos mapping of supraspinal and spinal micturition centers (Fig. 2J). In intact mice, repeated bladder filling induced robust c-Fos activation in the PAG and PMC. By contrast, SCI markedly reduced bladder filling–evoked activation within these supraspinal regions (Fig. 2K–N). Quantification confirmed significantly fewer c-Fos^+^ neurons in multiple PAG subregions, including the ventrolateral and lateral PAG, as well as in the PMC, in SCI mice relative to intact controls (Fig. 2M; Fig. S4A-B). In contrast, bladder filling induced comparable c-Fos activation in sacral regions, including the dorsal gray commissure, medial lateral horn, and dorsal horn in intact and SCI mice (Fig. 2N). Thus, bladder afferent signals continue to engage sacral spinal circuits after SCI but fail to effectively propagate to supraspinal micturition centers.

### Residual brain–bladder connectivity persists after SCI and remains functionally responsive

Previous studies show that neurons in the spared interlesion segment relay descending supraspinal signals to spinal circuits caudal to the injury^10,15^. In our prior work, staggered SCI induced prolonged swelling and progressive loss of excitatory neurons within this same region, contributing to functional dormancy of spared descending pathways^11^. We therefore asked whether residual brain–bladder communication after SCI depends on vulnerable relay neurons preserved between the two lesion sites.

To determine whether polysynaptic brain–bladder connectivity remains detectable after SCI, we injected the trans-synaptic retrograde tracer pseudorabies virus 152 (PRV-152), which encodes EGFP, into the bladder wall^19^ (Fig. 3A). Robust PRV-labeled neurons were observed in the pontine micturition center (PMC) of intact mice, whereas SCI markedly reduced, but did not eliminate, PMC labeling (Fig. 3B,C). Because direct Crh^+^ PMC projections to sacral spinal levels were abolished after staggered SCI (Fig. 2D), persistent PRV labeling in the PMC indicates that residual polysynaptic brain–bladder connectivity is not completely lost after injury.

**Figure 3.**
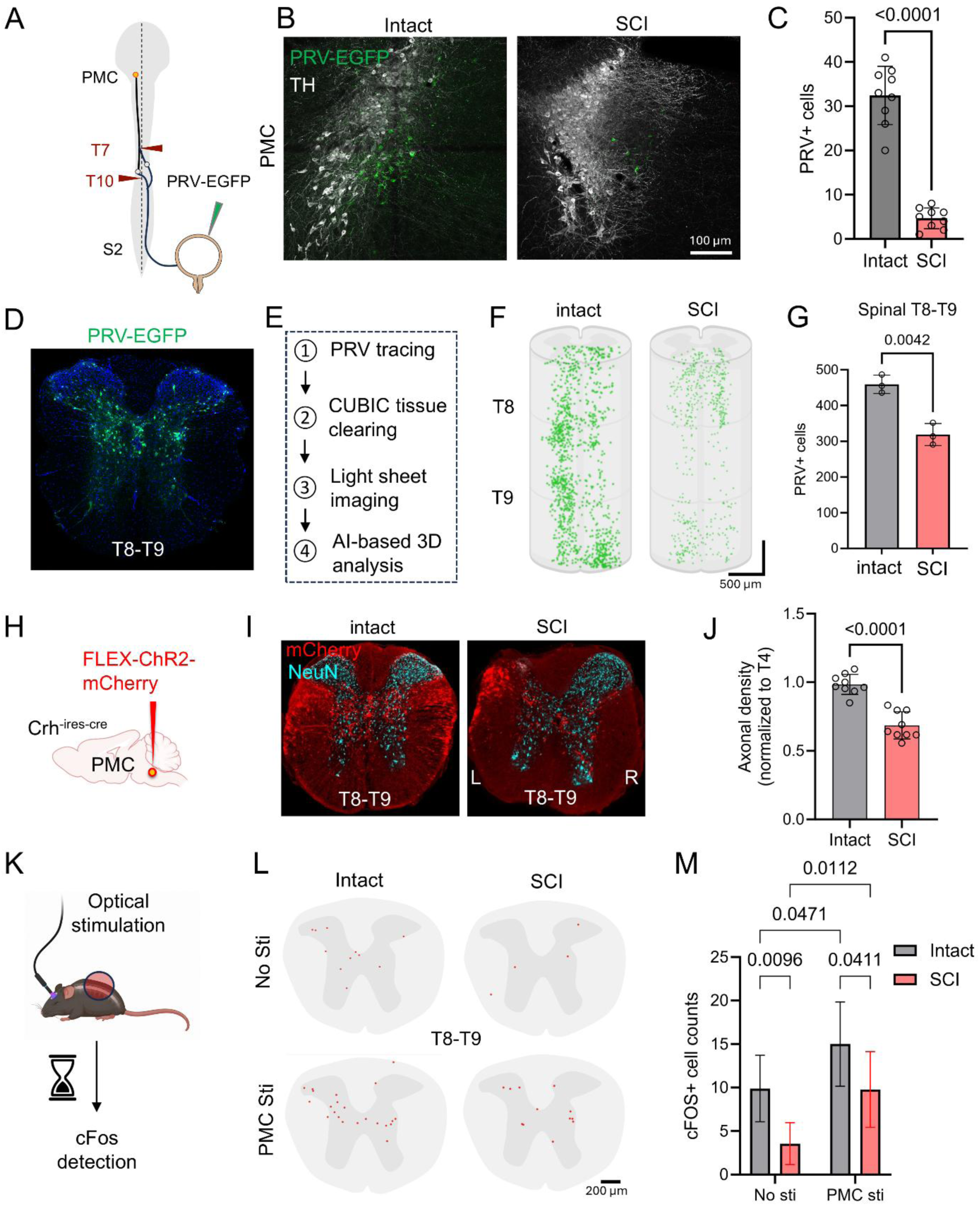
Anatomical and functional assessment of residual brain–bladder connectivity after SCI. (A) Schematic of PRV-152 retrograde trans-synaptic tracing from the bladder. (B, C) Representative PRV-152 labeling and quantification of PRV^+^ neurons in the pontine micturition center (PMC) of intact and SCI mice. N = 9, Unpaired t test with Welch’s correction. (D) Representative whole-mount PRV-152 labeling in the T8–T9 spinal segment. (E) Workflow for tissue clearing, light-sheet imaging, and machine learning-assisted three-dimensional image analysis. (F, G) Three-dimensional reconstruction and quantification of PRV^+^ neurons in the T8– T9 spinal segment of intact and SCI mice. N = 3, Unpaired t test with Welch’s correction. (H–J) Representative tracing and quantification of PMC-derived axons in the T8–T9 spinal segment of intact and SCI mice. N = 9, Unpaired t test with Welch’s correction. (K) Schematic of optogenetic stimulation of PMC Crh^+^ neurons followed by spinal c-Fos analysis. (L, M) Representative c-Fos staining and quantification of c-Fos^+^ neurons in the T8–T9 spinal segment following PMC stimulation in intact and SCI mice. N = 4-6, Two-way ANOVA followed by Tukey’s multiple comparisons test. Data are presented as mean ± SD. Scale bars are indicated in the panels.

We next asked whether bladder-connected neurons remained detectable within the T8–T9 interlesion region. To comprehensively quantify PRV-labeled neurons throughout this segment, we performed tissue clearing followed by whole-mount light-sheet imaging and AI-based three-dimensional cell segmentation (Fig. 3D–F). This approach enabled systematic three-dimensional reconstruction and quantification of PRV-positive neurons across the entire T8–T9 interlesion segment^11,20^. Although the number of PRV-labeled neurons was significantly reduced after SCI, a substantial population remained detectable within the preserved interlesion region (Fig. 3G), indicating that bladder-connected spinal neurons persist despite interruption of direct descending projections. Anatomical analyses further showed that spared PMC-derived mCherry-labeled axons traversed the interlesion bridge and extended into the T8–T9 region (Fig. 3H–J, Fig. S5A), while surviving neurons within this segment remained associated with descending serotonergic fibers (Fig. S5B), together indicating that the spared interlesion region retains anatomical features compatible with residual supraspinal brain–bladder communication.

To determine whether this preserved pathway remained functionally responsive to descending PMC input, we expressed ChR2 selectively in PMC Crh neurons and mapped spinal c-Fos activation following optical stimulation (Fig. 3K). In intact mice, PMC stimulation robustly activated neurons within the T8–T9 interlesion segment, whereas SCI significantly reduced, but did not abolish, this response (Fig. 3L,M). These findings demonstrate that residual supraspinal signals continue to engage neurons within the preserved T8–T9 interlesion network after SCI.

### Intrathecal bumetanide treatment improves urinary function after SCI

The persistence of residual anatomical connectivity and functional responsiveness within the brain–bladder network suggested that spared spinal circuitry might provide a substrate for therapeutic reinforcement. We therefore asked whether targeting dysfunction within the spared interlesion network could restore urinary function after SCI. Based on our previous finding that bumetanide reduces injury-induced neuronal swelling and restores conduction through spared spinal pathways^11^, staggered SCI mice received intrathecal bumetanide or vehicle continuously for 4 weeks beginning at the time of injury (Fig. 4A). In a 2-h VSOP assay, vehicle-treated mice produced sparse, small urine spots, whereas bumetanide-treated mice generated larger and more consolidated voids (Fig. 4B, C; Fig. S6), indicating improved voiding output. In the 8-h VSOP assay, vehicle-treated mice predominantly produced small, scattered droplets, whereas bumetanide-treated mice generated larger and more consolidated spots (Fig. 4D; Fig. S6; S7A-C). Leakage index showed a trend toward reduction but did not reach statistical significance (Fig. 4E; *p = 0.0575*). Gross morphology and quantification of residual urine volume further supported functional improvement: vehicle-treated SCI mice exhibited marked bladder distension, whereas bumetanide-treated mice showed reduced bladder enlargement (Fig. 4F, G), consistent with improved bladder emptying efficiency.

**Figure 4.**
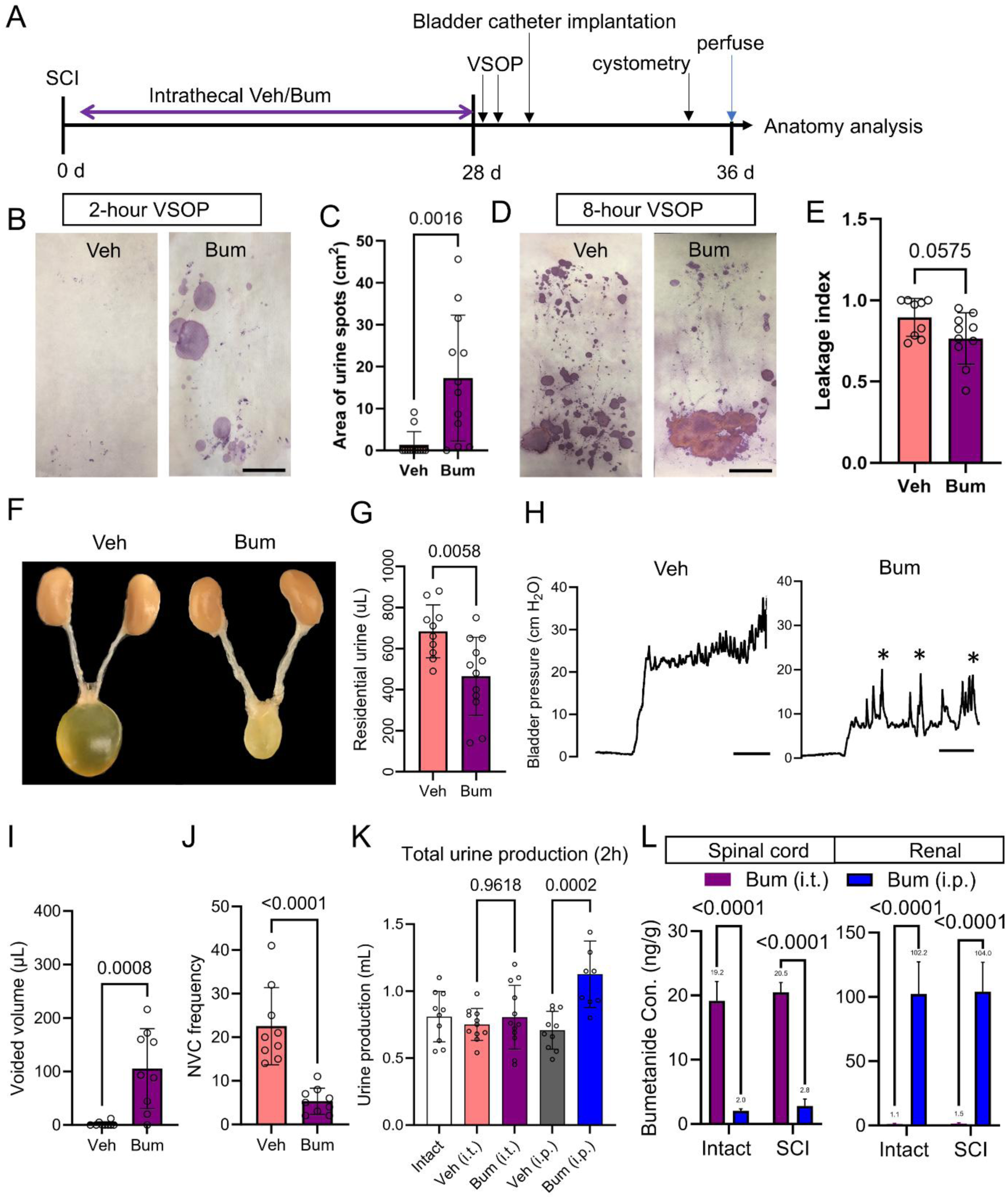
Intrathecal bumetanide improves bladder function after SCI. (A) Experimental timeline for SCI, continuous intrathecal vehicle (Veh) or bumetanide (Bum) treatment, VSOP testing, bladder catheter implantation, conscious cystometry, perfusion, and anatomical analysis. (B, C) Representative 2-h VSOP images and quantification of urine spot area in vehicle- and bumetanide-treated SCI mice. (D, E) Representative 8-h VSOP images and quantification of leakage index. N = 9-10, Unpaired t test with Welch’s correction. (F, G) Representative gross urinary tract morphology and quantification of residual urine volume. N = 10-12, Unpaired t test with Welch’s correction. (H) Representative conscious cystometric recordings from vehicle- and bumetanide-treated SCI mice. Asterisks indicate voiding contractions. (I, J) Quantification of voided volume and non-voiding contraction (NVC) frequency during conscious cystometry. N = 9, Unpaired t test with Welch’s correction. (K) Total urine production over 2 h in intact mice and SCI mice receiving vehicle or bumetanide by intrathecal (i.t.) or intraperitoneal (i.p.) administration. N = 10-12, One-way ANOVA followed by Tukey’s multiple comparisons test. (L) Bumetanide concentrations in the spinal cord and renal of intact and SCI mice following intrathecal or intraperitoneal administration. N = 6-8, Two-way ANOVA followed by Tukey’s multiple comparisons test. Data are presented as mean ± SD. Scale bars are indicated in the panels.

Conscious cystometry further demonstrated improved bladder dynamics following bumetanide treatment. Representative recordings showed sustained high-pressure filling with frequent non-voiding contractions in vehicle-treated SCI mice, whereas bumetanide-treated mice exhibited more discrete voiding contractions and reduced non-voiding activity (Fig. 4H). Quantitatively, bumetanide markedly increased voided volume and reduced the frequency of non-voiding contractions (Fig. 4I,J). Consistent with these improvements, bumetanide reduced residual urine volume, functional bladder capacity, and mean filling pressure, while increasing confirmed voiding frequency (Fig. S7D–G). Together, these findings demonstrate that intrathecal bumetanide improves both bladder emptying and storage dynamics after SCI.

Bumetanide inhibits renal NKCC2 and can cause diuresis, raising the possibility that its urinary benefits after SCI are secondary to increased urine production^13,14^. However, in intact mice, intrathecal (IT) bumetanide did not alter urine production in the 2-h assay, whereas systemic intraperitoneal (i.p.) administration increased urine output (Fig. 4K). Mass spectrometry further showed that i.p. delivery produced high renal but negligible spinal cord drug levels, whereas IT delivery enriched bumetanide in the spinal cord with minimal renal exposure (Fig. 4L), consistent with the low blood-brain barrier penetration rate of bumetanide^21^. These findings indicate that the therapeutic effects of intrathecal bumetanide are unlikely to result from diuresis and instead support a predominantly spinal mechanism that is distinct from increased urine production.

### Bumetanide strengthens bidirectional brain–bladder communication through the spared interlesion network

We next tested whether bumetanide enhanced bidirectional brain–bladder communication after SCI. Optogenetic stimulation of PMC Crh^+^ neurons rarely evoked bladder contractions in vehicle-treated SCI mice, whereas bumetanide increased both the proportion of stimulation trials eliciting a bladder response and the amplitude of evoked contractions (Fig. 5A–E), indicating improved descending supraspinal control. Conversely, bladder filling induced greater c-Fos activation in the PAG after bumetanide treatment, while c-Fos activation in the sacral spinal cord was unchanged between groups (Fig. 5F–J). Thus, bumetanide enhanced transmission across the injured spinal cord in both the descending PMC-to-bladder and ascending bladder-to-brain directions.

**Figure 5.**
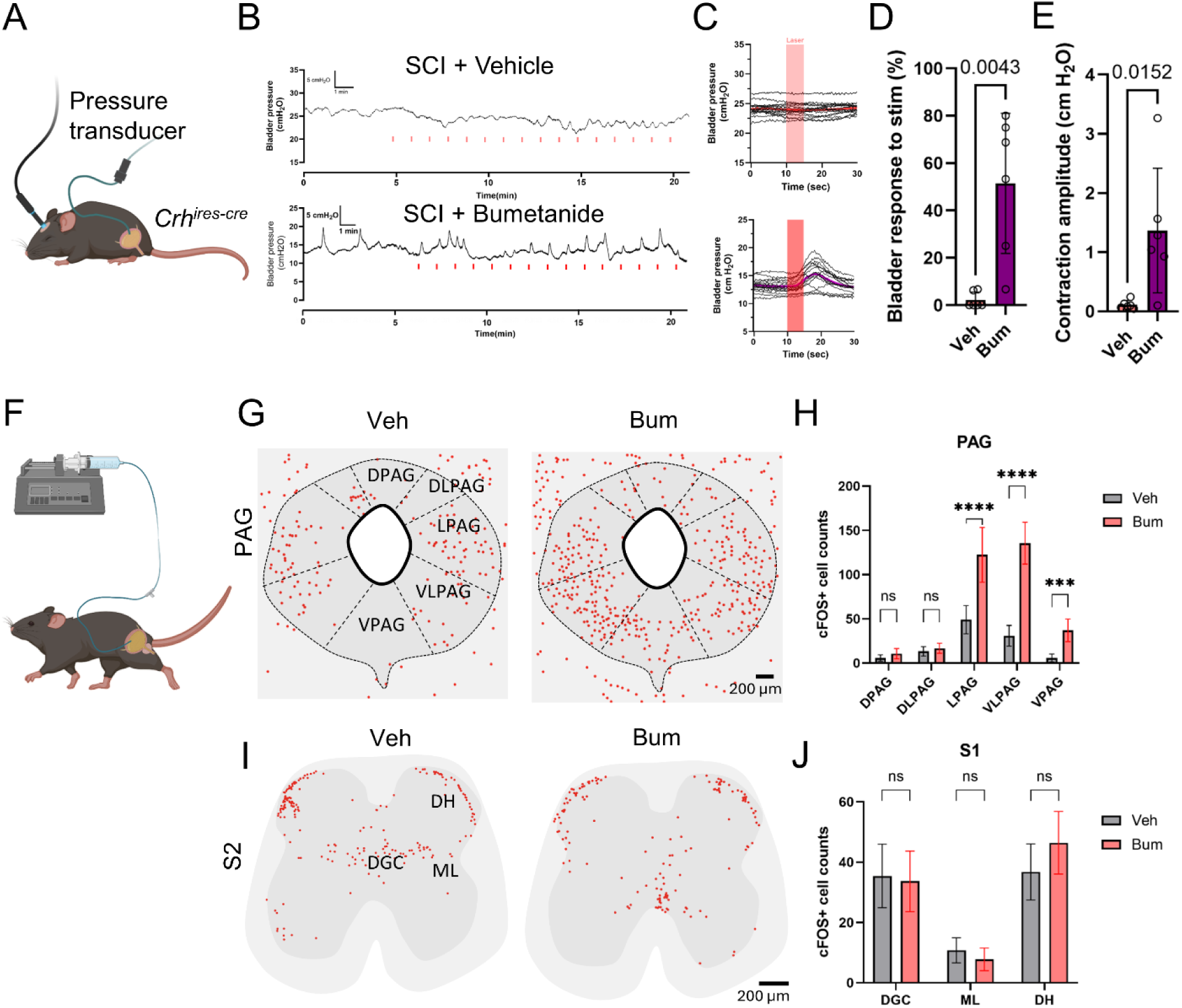
Intrathecal bumetanide strengthens bidirectional brain–bladder communication after SCI. (A) Schematic of non-invasive optogenetic activation of CRH+ PMC neurons and bladder pressure recording setup. (B) Representative bladder pressure traces during laser stimulation in vehicle- and bumetanide-treated SCI mice. (C) Bladder pressure aligned to laser onset of a representative mouse. (D–E) Quantification of bladder contraction response rate (D) and peak contraction amplitude (E) upon stimulation. N = 6, Unpaired t test with Welch’s correction. (F) Diagram of bladder infusion in awake mice and ascending bladder afferent pathway to S2 and PAG. (G) Representative c-Fos mapping in PAG regions after bladder filling in vehicle- and bumetanide-treated SCI mice. (H) Quantification of c-Fos+ neurons in PAG subregions (DPAG, LPAG, VLPAG, VPAG). N = 6, two-way ANOVA followed by Bonferroni multiple comparisons. (I) c-Fos+ neurons in the dorsal horn (DH) and dorsal gray commissure (DGC) at S2 spinal cord. (J) Quantification of c-Fos+ neurons in DH and DGC regions. N = 6, two-way ANOVA followed by Bonferroni multiple comparisons. Scale bars, 100 μm (G, I).

We next asked whether bumetanide-enhanced brain–bladder communication was associated with greater engagement of bladder-connected neurons within the spared T8–T9 interlesion region. Following bladder-wall injection of PRV-152, we cleared the T8–T9 spinal segment and performed whole-mount light-sheet imaging followed by AI-based three-dimensional cell detection and quantification (Fig. 6A,B). Vehicle-treated SCI mice retained a population of PRV-labeled neurons within T8–T9, consistent with the residual polysynaptic connectivity identified above. Bumetanide significantly increased the number of PRV^+^ neurons throughout this interlesion region (Fig. 6B,C). In contrast, PRV labeling within the L6–S2 spinal cord, including the spinal parasympathetic nucleus (SPN) and dorsal gray commissure (DGC)^6,7,16,22^, was comparable between vehicle- and bumetanide-treated mice (Fig. S8A), supporting consistent bladder PRV delivery and initial labeling across treatment conditions.

**Figure 6.**
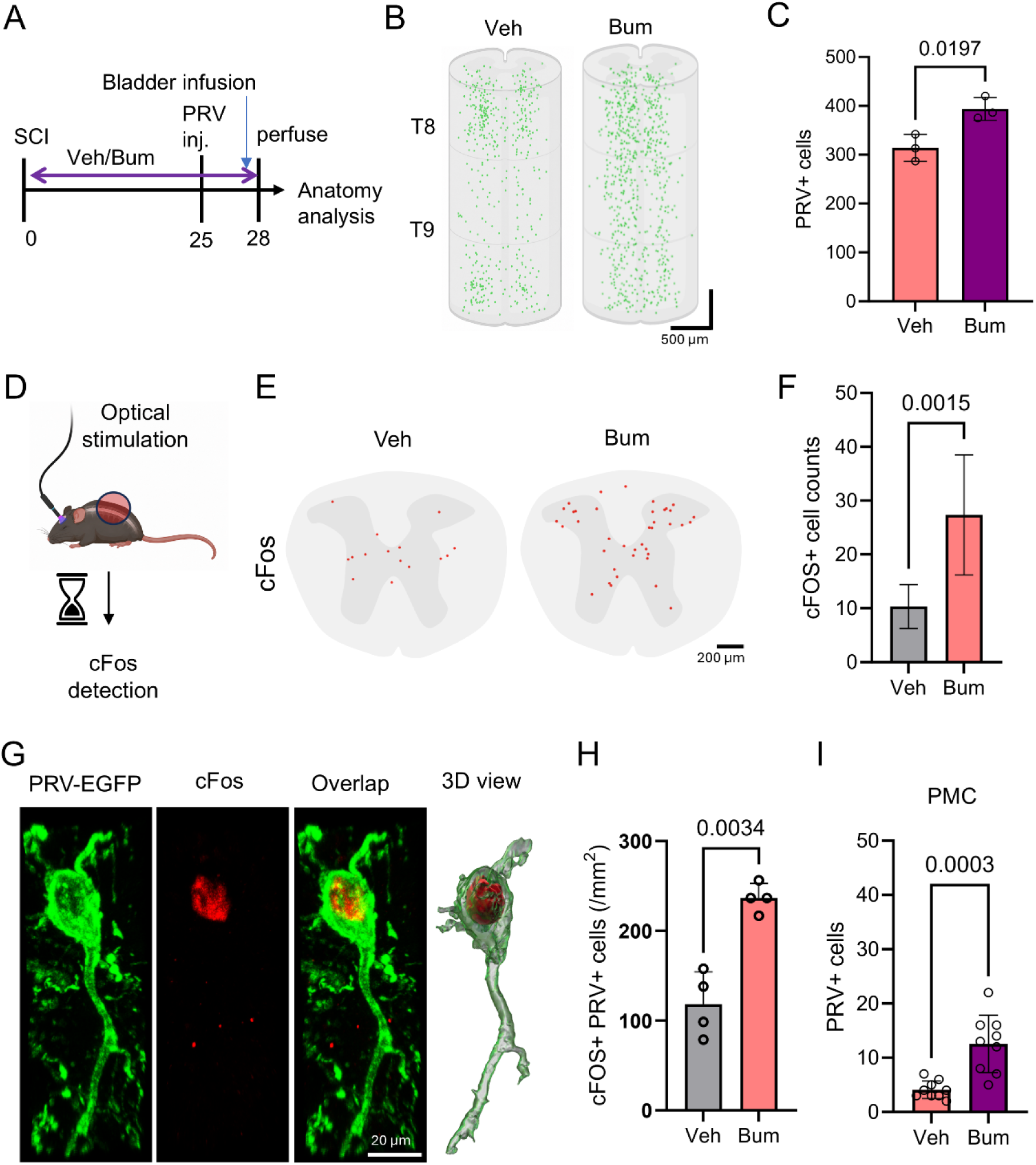
Bumetanide enhances functional engagement of spared interlesion neurons and brain–bladder connectivity after SCI. (A) Experimental timeline for SCI, continuous intrathecal vehicle (Veh) or bumetanide (Bum) treatment, bladder PRV-152 injection, perfusion, and anatomical analysis. (B, C) Representative three-dimensional reconstructions and quantification of PRV^+^ neurons within the T8–T9 interlesion spinal segment in vehicle- and bumetanide-treated SCI mice. N = 3, Unpaired t test with Welch’s correction. (D) Schematic of optogenetic stimulation followed by c-Fos detection in the spinal cord. (E, F) Representative c-Fos maps and quantification of c-Fos^+^ neurons within the T8–T9 interlesion segment in vehicle- and bumetanide-treated SCI mice. N = 6, Unpaired t test with Welch’s correction. (G) Representative images of PRV-EGFP^+^ neurons, c-Fos immunoreactivity, merged labeling, and three-dimensional reconstruction of a PRV^+^/c-Fos^+^ neuron. (H) Quantification of PRV^+^/c-Fos^+^ neurons within the T8–T9 interlesion segment. N = 4, Unpaired t test with Welch’s correction. (I) Quantification of PRV^+^ neurons in the pontine micturition center (PMC) following bladder PRV-152 injection. N = 9, Unpaired t test with Welch’s correction. Data are presented as mean ± SD. Scale bars, 500 μm (B), 200 μm (E), and 20 μm (G).

We next asked whether these bladder-connected interlesion neurons were responsive to descending PMC input. Following optogenetic stimulation of PMC Crh^+^ neurons, bumetanide-treated SCI mice showed significantly more c-Fos^+^ neurons within the T8–T9 interlesion region than vehicle-treated mice (Fig. 6D–F). Notably, PRV^+^/c-Fos^+^ neurons were also significantly increased after bumetanide treatment (Fig. 6G,H; Fig. S8B), identifying a larger population of bladder-connected interlesion neurons responsive to descending PMC output. Consistent with enhanced functional engagement, PRV labeling in the PMC was significantly increased after bumetanide treatment (Fig. 6I; Fig. S8C), consistent with enhanced polysynaptic brain–bladder connectivity. Altogether, these findings demonstrate that bumetanide strengthens bidirectional brain–bladder communication after SCI and enhances the functional engagement of spared T8– T9 interlesion neurons within this residual network.

## DISCUSSION

Neurogenic bladder is among the most disabling long-term sequelae after spinal cord injury (SCI), yet therapies that restore disrupted brain–bladder circuitry remain lacking^2,23^. Here, we show that SCI functionally uncouples both descending and ascending limbs of the spinobulbospinal micturition reflex despite evidence of residual anatomical connectivity. We identify the spared T8– T9 interlesion network as a candidate substrate for this residual communication and show that intrathecal bumetanide improves urinary function, strengthens bidirectional brain–bladder signaling, and enhances the functional engagement of spared interlesion neurons. These findings support a therapeutic strategy aimed at reinforcing residual spinal circuitry rather than requiring de novo circuit reconstruction.

Although modest in size, the human cohort provides an important clinical anchor for the study. AUA and NBSS profiles demonstrated persistent urinary incontinence, storage/voiding dysfunction, and impaired bladder-related quality of life, while increased post-void residual volume and representative urodynamic abnormalities were consistent with impaired emptying and abnormal bladder storage after SCI^2,24,25^. These clinical features were reproduced in the staggered SCI model, which developed urinary retention, fragmented voiding, leakage, increased residual urine, reduced voided volume, and frequent non-voiding contractions. Together, the human and experimental data establish a clinically relevant framework for interrogating the neural basis of urinary dysfunction after SCI.

A major conceptual advance of this study is that it reframes neurogenic bladder after SCI as a disorder of bidirectional brain–bladder communication rather than solely a consequence of peripheral bladder dysfunction. Classical models have emphasized loss of supraspinal descending control, detrusor overactivity, and detrusor–sphincter dyssynergia following SCI^2,5^.

Consistent with this view, staggered SCI markedly impaired PMC-evoked bladder contractions. However, our data also revealed disruption of the ascending limb of the circuit: bladder filling continued to activate sacral spinal regions but failed to effectively recruit supraspinal PAG–PMC centers. Because the PAG integrates bladder afferent information and gates voiding through interactions with the PMC^4,26,27^, these findings indicate that SCI disrupts signal transmission in both directions across the injured spinal cord. Importantly, bumetanide enhanced PMC-evoked bladder responses while also increasing bladder filling–evoked PAG activation without altering sacral spinal activation, supporting improved transmission of bladder-derived signals to supraspinal centers rather than a nonspecific increase in peripheral sensory activation. Thus, the therapeutic effect of bumetanide appears to involve enhancement of bidirectional communication rather than isolated enhancement of bladder contractility.

Our findings further extend the concept of spared relay-mediated recovery after SCI from locomotor to autonomic function. In incomplete SCI, residual supraspinal signals can be rerouted through propriospinal or intraspinal interneuronal networks to support locomotor recovery^10,15^. Lower urinary tract control similarly depends on distributed spinal interneuronal circuits, and SCI-induced plasticity within these networks can contribute to both adaptive reflex recovery and maladaptive bladder dysfunction^8,28^. In the present study, bladder-connected PRV^+^ neurons persisted within the T8–T9 interlesion region after SCI, PMC-derived axons remained detectable within this segment, and PMC stimulation continued to activate T8–T9 neurons. These complementary anatomical and functional observations identify the spared T8–T9 network as a candidate substrate through which residual supraspinal signals may engage bladder-related spinal circuitry after SCI.

The treatment experiments further support this interpretation. Whole-mount light-sheet imaging and AI-based three-dimensional quantification demonstrated increased PRV labeling within T8– T9 after bumetanide treatment, while PRV labeling within the L6–S2 bladder-related spinal circuitry remained comparable between treatment groups, supporting consistent initial PRV delivery and labeling. Bumetanide also increased PMC stimulation–evoked c-Fos activation within T8–T9 and, importantly, increased PRV^+^/c-Fos^+^ neurons, identifying a larger population of bladder-connected interlesion neurons that remained responsive to descending supraspinal input. This enhanced functional engagement was accompanied by increased PRV labeling in the PMC, consistent with strengthened long-range connectivity between the bladder and brain. Rather than generating a de novo pathway, these findings suggest that bumetanide acts on a pre-existing residual network that remains anatomically present but functionally compromised after SCI.

Our previous work provides a potential mechanistic basis for this effect. We previously showed that intrathecal bumetanide reduces prolonged neuronal swelling and preserves interlesion neurons after staggered SCI^11^. NKCC1 contributes to ionic dysregulation, edema, and excitotoxic vulnerability after central nervous system injury^29–31^, and persistent swelling within a narrow interlesion region could compromise signal propagation even when anatomical continuity is retained. The present study does not directly quantify neuronal preservation after bumetanide treatment, but the increased PRV labeling, PMC-evoked c-Fos responses, and PRV^+^/c-Fos^+^ neuronal population observed within T8–T9 are consistent with greater functional engagement of the spared interlesion network. One possibility is therefore that the neuroprotective effect identified in our previous study preserves the structural substrate necessary for relay transmission, allowing residual brain–bladder signals to propagate more effectively. Additional effects on spinal excitability, chloride homeostasis, or inhibitory–excitatory balance may also contribute and will require further investigation.

These findings have important translational implications for neurogenic bladder after SCI. Current management—including intermittent catheterization, antimuscarinic therapy, botulinum toxin, and surgical or device-based interventions—can reduce complications and improve bladder storage or emptying but does not directly restore supraspinal control of micturition^2^. In contrast, our data suggest that intrathecal bumetanide acts at the level of residual spinal circuitry. Intrathecal delivery produced spinal exposure without increasing overall urine production, supporting a spinal rather than diuretic mechanism of action. This distinction is important because systemic bumetanide is a potent loop diuretic with limited CNS penetration and dose-limiting peripheral effects^12–14^. Although intrathecal drug delivery is clinically established for selected neurological indications such as severe spasticity^32^, intrathecal bumetanide for SCI will require dedicated studies of dosing, safety, and therapeutic window. More broadly, pharmacologic reinforcement of spared circuitry may complement regenerative and activity-based approaches. Epidural stimulation and related neuromodulatory strategies can uncover residual motor and autonomic pathways after chronic SCI^33–37^, and preserving or strengthening the spinal substrate available for re-engagement may increase the effectiveness of such interventions.

Several limitations warrant consideration. First, although the staggered hemisection model provides experimental access to spared interlesion circuitry, it does not capture the anatomical and clinical heterogeneity of human contusive SCI. Whether similar residual relay mechanisms operate across other injury types remains to be determined. Second, our convergent anatomical and functional data nominate the T8–T9 interlesion region as a candidate relay substrate, but direct manipulation of defined neuronal populations will be required to establish necessity and sufficiency. Identifying whether specific excitatory, inhibitory, or genetically defined propriospinal populations mediate residual or recovered signaling will be an important next step. Third, bumetanide was initiated at the time of injury; whether delayed treatment can restore established chronic urinary dysfunction remains unknown. Finally, studies defining the optimal intrathecal dose, therapeutic window, long-term safety, and compatibility with existing bladder-management strategies will be necessary before clinical translation.

In summary, our study reframes neurogenic bladder after SCI as a disorder of bidirectional communication within a partially preserved autonomic network. We identify the spared T8–T9 interlesion network as a candidate substrate for residual brain–bladder signaling and show that intrathecal bumetanide strengthens bidirectional communication while enhancing the functional engagement of spared interlesion neurons. These findings support therapeutic reinforcement of residual spinal circuitry as a potential strategy for promoting autonomic recovery after SCI.

## STAR METHODS

KEY RESOURCES TABLE.

KEY RESOURCES TABLE
| REAGENT OR RESOURCE | SOURCE | IDENTIFIER |
| --- | --- | --- |
| Experimental models: Organisms/strains |  |  |
| Wild-type mice | The Jackson Laboratory | #000664 |
| Crh<sup>IRES-Cre</sup> mice | The Jackson Laboratory | #012704 |
| Ai14 reporter mice (Rosa26-loxP-STOP-loxP-tdTomato) | The Jackson Laboratory | #007914 |
| Crh<sup>IRES-Cre</sup>::Ai14 mice | Generated by intercrossing in this study | N/A |
| Viral strains |  |  |
| AAV2/8-Flex-ChR2-mCherry | Boston Children’s Hospital Viral Core | NA |
| AAV2/8-hSyn-ChR2-eYFP | Salk Institute | NA |
| AAV-nEF-Con/Foff2.0-ChRmine-oScarlet | Salk Institute | NA |
| AAV-nEF-Con/Foff2.0-oScarlet | Salk Institute | NA |
| PRV-152 (GFP-expressing pseudorabies virus) | Gift from Zhang laboratory, UT Southwestern Medical Center | NA |
| Antibodies |  |  |
| Anti-GFAP | Invitrogen | #13-0300; 1:600 |
| Anti-TH | Millipore | #152; 1:2,000 |
| Anti-NeuN | Millipore | #ABN90P; 1:1,000 |
| Anti-GFP | Abcam | #ab13970; 1:500 |
| Anti-c-Fos | Synaptic Systems | #226308; 1:500 |
| Anti-5-HT | Thermo Fisher Scientific | #BS-1126R; 1:400 |
| Alexa Fluor 488/555 secondary antibodies | Thermo Fisher Scientific | #A-21206, A-31572, A-21208, A78948; 1:300 |
| Alexa Fluor 647 secondary antibodies | Jackson Immuno | #706-605-148 |
| Chemicals, peptides, and recombinant proteins |  |  |
| Bumetanide | Tocris | #3108 |
| Artificial cerebrospinal fluid (aCSF) | Tocris | #3525 |
| Ninhydrin | Sigma-Aldrich | #151173 |
| Fluoromount-G | Invitrogen | #00-4959-52 |
| CUBIC-L | TCI America | #T3740 |
| CUBIC-R+(M) | TCI America | #T3741 |
| Critical commercial assays/equipment |  |  |
| ALZET osmotic pump | ALZET | #1004 |
| Intrathecal catheter | ALZET | #0007743 |
| Small Animal Cystometry Lab Station | Med Associates | NA |
| Microinjector | World Precision Instruments | Nanoliter 2000 |
| Lightsheet microscope | Zeiss | Z.1 |
| Confocal microscope | Zeiss | LSM880 |
| Nikon Eclipse Ti2 fluorescence microscope | Nikon | Eclipse Ti2 |
| Software and algorithms |  |  |
| GraphPad Prism | GraphPad Software | Version 10 |
| Fiji/ImageJ | NIH | NA |
| Imaris | Oxford Instruments | Version 10.0.1 |
| Imaris File Converter | Oxford Instruments | Version 10.0.1 |
| Labkit | Fiji/ImageJ plugin | NA |
| SOF-552 cystometry analysis software | Catamount Research and Development | SOF-552 |
| MATLAB | MathWorks | 2024b |
| Python | Python.org | v3.13 |

## RESOURCE AVAILABILITY

### Lead contact

Further information and requests for resources and reagents should be directed to and will be fulfilled by the lead contacts, Bo Chen and Qiang Li.

### Materials availability

This study did not generate new unique reagents.

### Data and code availability

All data reported in this paper will be shared by the lead contacts upon reasonable request. Custom code used for data analysis and visualization is publicly available at GitHub: https://github.com/qiali2020/sci-neurogenic-bladder

## EXPERIMENTAL MODEL AND STUDY PARTICIPANT DETAILS

### Human participants

Seventeen individuals with chronic spinal cord injury (SCI) were prospectively enrolled between 2021 and 2025 under approval from the University of Texas Medical Branch (UTMB Health) Institutional Review Board (IRB protocol no. 20-0197), with written informed consent obtained from all participants. Demographic and clinical characteristics, including age, sex, neurological level of injury, injury completeness, injury etiology, and time since injury, were recorded. All participants used clean intermittent catheterization for bladder management.

### Mice

Adult male and female wild-type mice (The Jackson Laboratory [JAX] #000664), Crh^IRES-Cre^ mice^38^ (JAX #012704), Ai14 reporter mice^39^ (Rosa26-loxP-STOP-loxP-tdTomato; JAX #007914), and Crh^IRES-Cre^::Ai14 mice generated by intercrossing these lines were used. Mice were 8–12 weeks old at the time of experiments and housed under standard conditions with ad libitum access to food and water on a 12-h light/dark cycle. Animals were randomly assigned to experimental groups, and investigators were blinded during data collection and analysis when feasible. All animal procedures were approved by the University of Texas Medical Branch Institutional Animal Care and Use Committee (protocol #1910082) and were performed in accordance with institutional guidelines.

## METHOD DETAILS

### Human cohort and urinary function assessment

Neurogenic lower urinary tract symptoms were assessed using the Neurogenic Bladder Symptom Score (NBSS), which evaluates urinary incontinence, storage and voiding symptoms, complications, and bladder-related quality of life. Lower urinary tract symptom burden was additionally assessed using the American Urological Association Symptom Index (AUA-SI), including the associated quality-of-life assessment. Individual patient scores were used to characterize the severity and distribution of urinary symptoms across the cohort. Objective bladder dysfunction was assessed using post-void residual (PVR) urine volume and urinary leakage frequency. PVR measurements were obtained as part of the clinical evaluation. Urinary leakage was recorded using a 5-day bladder diary, and the average number of leakage episodes per day was calculated for each participant.

Multichannel urodynamic studies were performed as part of the clinical evaluation using standard clinical procedures. Intravesical pressure was monitored during bladder filling and voiding to assess abnormalities in bladder storage and emptying. A representative urodynamic tracing was included to illustrate characteristic dysfunction observed in chronic SCI, including elevated bladder pressure during filling, urinary leakage preceding coordinated voiding, and incomplete bladder emptying. Because complete quantitative urodynamic datasets were not available for all participants, urodynamic variables were not included in cohort-level statistical analyses. Clinical data were summarized at the individual-patient and cohort levels. NBSS domain scores, AUA-SI scores, PVR volumes, and average daily leakage episodes were used to define the major clinical features of neurogenic bladder dysfunction subsequently evaluated in the experimental SCI model.

### Staggered spinal cord injury

For surgical procedures, mice were anesthetized with ketamine/xylazine (100/10 mg/kg, intraperitoneal) and received sustained-release buprenorphine (0.1 mg/kg, subcutaneous) for perioperative analgesia. Following a midline dorsal incision, laminectomies were performed at T7 and T10. A right-sided over-hemisection was made at T7 using a 15° Feather microscalpel (Electron Microscopy Sciences #7204515) and micro-scissors (Fisher Scientific #12-460-288), interrupting the dorsal columns and ipsilateral ventrolateral pathways without sparing. A complementary left-sided hemisection was then created at T10, extending to the spinal midline while preserving the contralateral side. This staggered bilateral lesion paradigm interrupts descending and ascending pathways while preserving an interlesion spinal segment, as previously described^9^. Muscle layers were sutured and the skin was closed with wound clips. Animals were maintained on a heating pad until recovery from anesthesia.

### Post-surgical treatment and animal care

At the completion of surgical procedures, mice received 1 mL sterile saline subcutaneously to reduce the risk of dehydration. Animals were monitored twice daily for general activity, wound healing, aggressive interactions, biting wounds, and signs of urinary infection or other complications. Manual bladder expression was performed twice daily as needed. If open wounds or signs of infection were detected, enrofloxacin (Baytril; 10 mg/kg) was administered for 7 days. Body weight was recorded weekly.

### Viral delivery for circuit mapping

For PMC Crh+ neuron labeling, adult Crh^IRES-Cre^ mice were anesthetized with ketamine/xylazine and placed in a stereotaxic frame (David Kopf Instruments) under aseptic conditions. A small burr hole was drilled, and AAV2/8-Flex-ChR-mCherry (Boston Children’s Hospital Viral Core; titer, 0.5 × 10^13^) was injected unilaterally into the pontine micturition center (PMC) through a pulled glass micropipette. A total volume of 150 nL was delivered at 100 nL/min at coordinates 5.25 mm posterior to bregma, 0.70 mm lateral, and 3.15 mm ventral from the pial surface. The pipette was left in place for 5 min after injection to minimize reflux before slow withdrawal.

For T8–T9 spinal viral delivery, adult wild-type mice underwent laminectomy over T8–T9 and received bilateral intraspinal injections of AAV2/8-hSyn-ChR2-eYFP (Salk Institute; titer, 1.7 × 10^12^) using a pulled glass micropipette beveled at a 25° angle. Injections were made 1 mm lateral to the midline at depths of 0.5 and 1.0 mm from the dorsal spinal surface. Three injection sites were spaced 1 mm apart along the rostrocaudal axis to target the interlesion segment (100 nL per site). Viral vectors were delivered using a Nanoliter 2000 microinjector (World Precision Instruments), and the pipette was left in place briefly after each injection to minimize reflux. Animals recovered for at least 3 weeks before downstream experiments.

### Non-invasive optogenetic stimulation of PMC Crh+ neurons

Non-invasive deep-brain optogenetic stimulation was performed as previously described, with modifications for PMC circuit interrogation (PMID: 33020604). Adult *Crh*^IRES-Cre^ mice were anesthetized with 2% isoflurane and placed in a stereotaxic apparatus on a thermostatically controlled heating pad. AAV-nEF-Con/Foff2.0-ChRmine-oScarlet (Salk Institute Vector Core; titer, 4.7 × 10^12^ vg/mL) or the corresponding control virus AAV-nEF-Con/Foff2.0-oScarlet (Salk Institute Vector Core; titer, 2 × 10^12^ vg/mL) was stereotaxically injected into the PMC using pulled glass micropipettes at coordinates AP −5.25 mm, ML ±0.70 mm, and DV −3.15 mm from the pial surface. A total volume of 200 nL per site was delivered at approximately 100 nL/min, and the injection pipette was maintained in place for 5 min before withdrawal. Mice were allowed to recover for 2 weeks to permit stable viral expression.

Before physiological experiments, a 200- or 400-μm optic fiber coupled to a stainless-steel ferrule (0.39 numerical aperture; Thorlabs) was positioned over the intact skull above the PMC target region and secured with Metabond dental cement. The ferrule was connected to a 635-nm laser source via a fiber-optic patch cable. Laser output was controlled by a pulse generator (CK10210, CNI Laser), with the laser controller set to 400 mW. PMC neurons were stimulated using 5-ms pulses at 20 Hz. Each stimulation epoch lasted 5 s and was repeated once per minute (5 s ON, 55 s OFF).

During bladder physiology experiments, optical stimulation was synchronized with conscious cystometry, and intravesical pressure was continuously recorded. Physiological responses were analyzed offline using custom MATLAB scripts. For neuronal activation experiments, the same stimulation paradigm was applied over a 5-min period, and mice were perfused 90 min after stimulation for c-Fos immunohistochemistry. Viral expression and targeting accuracy were confirmed histologically by oScarlet fluorescence within the PMC and its descending projections. Animals with mistargeted injections or insufficient viral expression were excluded from analysis.

### Pseudorabies virus bladder-wall tracing

Trans-synaptic retrograde tracing of brain–bladder circuitry was performed using PRV-152, a GFP-expressing pseudorabies virus (gift from the Zhang laboratory, UT Southwestern Medical Center; 7.9 × 10^8^ PFU/mL). Mice were anesthetized with ketamine/xylazine (100/10 mg/kg, intraperitoneal), and the bladder was exposed through a lower midline abdominal incision under aseptic conditions. PRV-152 was injected into two sites of the bladder wall at 500 nL per site using a pulled glass micropipette, for a total injection volume of 1 μL per animal, to achieve distributed uptake while avoiding leakage into the bladder lumen. The abdominal wall and skin were sutured in layers, and animals were monitored during recovery. Mice were maintained for 3 days after viral injection to permit trans-synaptic retrograde labeling, after which they were perfused for tissue collection. PRV-labeled neurons were quantified in the PMC and spinal cord regions associated with bladder control.

### Bumetanide preparation

Bumetanide (Tocris #3108) was prepared separately for intrathecal and systemic administration. For intrathecal infusion, bumetanide was dissolved in sterile artificial cerebrospinal fluid (aCSF; Tocris #3525) containing 10% DMSO and 1 mM NaOH at a final concentration of 0.37 mg/mL, corresponding to an estimated delivery dose of approximately 0.05 mg/kg/day. For intraperitoneal administration, bumetanide was formulated in sterile saline containing 10% DMSO and 1 mM NaOH at 0.02 mg/mL, yielding an approximate daily dose of 0.2 mg/kg. Vehicle solutions contained the corresponding solvent formulations without bumetanide. All solutions were sterile-filtered before use.

### Systemic bumetanide administration

For systemic administration, bumetanide was delivered by intraperitoneal injection once daily beginning on the day of SCI and continued for 4 weeks. Bumetanide was administered at approximately 0.2 mg/kg per day using the formulation described above. Vehicle-treated animals received an equivalent volume of vehicle on the same schedule.

### Continuous intrathecal drug delivery

For continuous intrathecal delivery of bumetanide or vehicle, osmotic pumps (100 μL capacity; ALZET model 1004) were connected to intrathecal catheters and prepared under aseptic conditions. Pumps were filled with bumetanide or vehicle using an ALZET filling needle, and the flow moderator was filled with the same solution. Filled pumps were primed in sterile saline at 37°C for 48 h before implantation.

For implantation, mice were anesthetized with ketamine/xylazine (100/10 mg/kg, intraperitoneal). A small incision was made over the lumbosacral spine, and the L6 vertebra was exposed. Following laminectomy, a small dural opening was made between L6 and S1 using a 30-gauge needle. An intrathecal catheter (ALZET #0007743) with a 3-mm trimmed tip was inserted rostrally into the intrathecal space with the aid of a Teflon-coated stylet. After removal of the stylet and confirmation of cerebrospinal fluid entry into the catheter, the catheter was secured with surgical glue and 7-0 sutures and connected to the osmotic pump through a metal connector. Pumps were implanted subcutaneously in the dorsal flank and delivered bumetanide or vehicle at 0.11 μL/h for the indicated experimental duration. Bumetanide or vehicle treatment was initiated on the day of injury, with osmotic pump implantation and intrathecal catheter placement performed approximately 0.5–1 h after completion of the staggered SCI surgery. Continuous intrathecal infusion was maintained for 4 weeks. Animals were monitored daily, and pumps were gently mobilized during routine care to minimize adhesion formation.

### Bladder catheter implantation and conscious cystometry

For bladder catheter implantation, mice were anesthetized with ketamine/xylazine (100/10 mg/kg, intraperitoneal), and a lower midline abdominal incision was made under sterile conditions. A saline-filled PE-10 catheter with a heat-flared tip (Med Associates) was inserted into the bladder dome and secured with a 6-0 nylon purse-string suture. Using a polished 26-gauge needle as a guide, the distal catheter was tunneled subcutaneously to the dorsal neck region, exteriorized, sealed, and temporarily positioned within the subcutaneous space. Muscle and skin layers were closed separately with non-absorbable sutures. Sustained-release buprenorphine (0.1 mg/kg, subcutaneous) was administered immediately after surgery and again 3 days later. Enrofloxacin (2.5 mg/kg in 1 mL saline, subcutaneous) was administered once daily during the postoperative recovery period. Animals recovered for 72 h before cystometric testing. Before cystometric recordings, mice were habituated to restrainers twice daily for 30-min sessions over two consecutive days. Conscious cystometry was performed using a Small Animal Cystometry Lab Station (Med Associates). Mice were placed in flat-bottom restrainers (Plas Labs) positioned above a wire grid and an electronic balance, permitting simultaneous recording of intravesical pressure and urine output. A sheet of Parafilm was placed on the balance to collect voided urine. To minimize adherence of urine droplets to the wire grid and facilitate passage of urine to the collection surface, the grid was lightly coated with WD-40 before recording. The balance was tared after the collection setup was assembled.

The bladder catheter was connected through a pressure transducer to a syringe pump and data-acquisition system. Room-temperature sterile saline was continuously infused at 0.01 mL/min. After a 20–30 min stabilization period, cystometric activity and urine output were recorded continuously for a 30-min analysis period, during which at least three reproducible filling–voiding cycles were obtained whenever possible. Urine released during the recording period was collected continuously on the Parafilm-covered balance. Cumulative voided volume was defined as the total volume of urine collected during the 30-min recording period. A confirmed voiding event was defined as visually observed urine release onto the collection surface, accompanied by a corresponding increase in the balance signal. Confirmed voiding frequency was defined as the total number of such events during the 30-min recording period. Cystometric recordings were analyzed in a blinded manner using SOF-552 software (Catamount Research and Development). Non-voiding contractions (NVCs) were defined as elevations in intravesical pressure >5 cm H₂O above the filling baseline without detectable urine release. Functional bladder capacity during continuous cystometry was calculated from the volume infused during the intermicturition interval. Filling pressure was assessed from the baseline intravesical pressure during the bladder-filling phase; when NVCs were present, nadir pressures between NVCs were used to estimate the filling baseline, consistent with recommended terminology for rodent cystometry^40^. Mean filling pressure was obtained by averaging filling-pressure measurements across analyzed cycles within each animal. At the end of the recording session, saline infusion was stopped and residual bladder volume was collected through the catheter and measured. Each mouse was treated as an independent biological replicate. For cystometric parameters measured on a cycle-by-cycle basis, values from at least three reproducible cycles were averaged to generate a single value for each animal before statistical analysis. Cumulative voided volume and confirmed voiding frequency were calculated over the complete 30-min recording period and therefore yielded one value per animal.

### Voiding spot on paper assay

Voiding behavior was assessed using the voiding spot on paper (VSOP) assay 4 weeks after SCI, as previously described. Mice were manually bladder-expressed immediately before testing to standardize baseline bladder volume and were then placed individually in standard mouse cages (34 × 16 cm) lined with Whatman filter paper covering the cage floor. VSOP assays were performed for 2-h and 8-h collection periods using fresh cages and filter paper for each test. All assays began at 9:00 a.m. During testing, mice were provided standard dry chow and gel hydration packs.

After collection, filter papers were stained with ninhydrin (Sigma-Aldrich) to visualize urine spots and digitally scanned. Images were imported into Fiji/ImageJ, calibrated, and analyzed by measuring individual urine spots using the freehand selection tool. Overlapping spots were counted as a single void unless clearly separable borders were visible. Voiding frequency was defined as the total number of discrete urine spots during the collection period. The leakage index was calculated as the proportion of small urine spots <0.5 cm in diameter relative to the total number of voids.

### Residual urine measurement

Residual urine volume was assessed after urinary behavioral testing. Mice were gently restrained, and the bladder was manually expressed by applying mild pressure to the lower abdomen until no additional urine could be expelled. Expelled urine was collected and quantified as residual urine volume. Bladder expression was performed consistently across experimental groups and time points.

### Bladder filling-induced c-Fos activation

Activity-dependent neuronal activation in bladder-control circuits was assessed by c-Fos induction following bladder filling. Mice were anesthetized, and a PE-10 catheter was inserted into the bladder dome and secured with a purse-string suture. The catheter was tunneled subcutaneously from the abdomen to the dorsal neck region, and the distal end was sealed using heated forceps and temporarily placed beneath the skin. After a 3-day recovery period, the catheter was exteriorized and connected to the cystometry system.

In awake mice, the bladder was continuously infused with sterile saline at 40 μL/min for 30 min to evoke bladder afferent signaling and micturition-related circuit activation. Control animals underwent the same catheterization and connection to the cystometry system but received no saline infusion. Following bladder stimulation, mice were maintained for up to 30 min before perfusion to allow c-Fos expression. Brain and spinal cord tissues were then processed for c-Fos immunofluorescence, and c-Fos-positive cells were quantified in bladder-control regions including the periaqueductal gray, pontine micturition center, and sacral spinal cord.

### Immunofluorescence

At the completion of experiments, mice were deeply anesthetized with ketamine/xylazine (100/10 mg/kg, intraperitoneal) and transcardially perfused with phosphate-buffered saline (PBS) followed by 4% paraformaldehyde (PFA) in PBS. Brain and spinal cord tissues were dissected and post-fixed overnight in 4% PFA at 4°C, then cryoprotected in 30% sucrose for at least 48 h. Samples were embedded in plastic molds (Thermo Fisher Scientific #22-19) and snap-frozen in dry ice-cooled ethanol. Spinal cord segments spanning C2–S2 were cut into continuous 1.5-mm tissue blocks and embedded with the rostral surface oriented downward to preserve rostrocaudal alignment. Frozen tissues were sectioned at 40 μm on a Leica CM1950 cryostat, and free-floating sections were collected.

Sections were blocked for 1 h at room temperature in PBS containing 10% donkey serum and 0.5% Triton X-100, then incubated overnight at 4°C with primary antibodies diluted in blocking buffer: Rat anti-GFAP (Invitrogen #13-0300, 1:600), Rabbit anti-TH (Millipore #152, 1:2,000), Guinea pig anti-NeuN (Millipore #ABN90P, 1:1,000), Chicken anti-GFP (Abcam #ab13970, 1:500), Guinea pig anti-c-Fos (Synaptic Systems #226308, 1:500), and Rabbit anti-5-HT (Thermo Fisher Scientific #BS-1126R, 1:400). After three PBS washes, sections were incubated with species-appropriate Alexa Fluor 488, 555, or 647 secondary antibodies (Thermo Fisher Scientific # A-21206, A-31572, A-21208, A78948; Jackson Immuno # 706-605-148) for 2 h at room temperature. Sections were washed, mounted on charged slides, and coverslipped with Fluoromount-G (Invitrogen #00-4959-52).

Fluorescence images were acquired using a Nikon Eclipse Ti2 inverted fluorescence microscope or a Zeiss LSM880 confocal laser-scanning microscope. Acquisition settings were held constant across experimental groups for quantitative comparisons. Brightness/contrast adjustment, pseudo-color assignment, and cropping were performed in ImageJ for presentation only. For quantitative analysis, 4–5 sections spanning the dorsal–ventral axis were imaged for each spinal cord region.

### Bladder histology

After transcardial perfusion, urinary bladders were dissected and fixed in 4% PFA at 4°C. Samples were submitted to the Clinical Pathology Laboratory at Jennie Sealy Hospital, University of Texas Medical Branch (UTMB), for paraffin embedding, sectioning at 5 μm, and automated hematoxylin and eosin staining using standard histological procedures. Stained bladder sections were imaged using an Olympus inverted light microscope under bright-field illumination.

### Mass spectrometric quantification of bumetanide

To assess route-dependent tissue exposure following drug administration, spinal cord and kidney tissues were harvested from animals receiving intrathecal or systemic bumetanide. Tissues were rapidly collected, weighed, and flash-frozen until processing. For drug extraction, samples were homogenized in acetonitrile at a ratio of 1 mL solvent per gram of tissue and centrifuged at 12,000 rpm for 10 min. The resulting supernatants were collected, and an aliquot from each sample was submitted to the University of Texas Medical Branch Mass Spectrometry Facility (UTMB MSF) for quantitative measurement of bumetanide by liquid chromatography–mass spectrometry. Bumetanide concentrations were normalized to tissue weight and compared between spinal cord and kidney samples following intrathecal versus systemic administration.

### Tissue clearing and light-sheet imaging

For tissue clearing and three-dimensional imaging, spinal cord samples were processed using a modified CUBIC-based protocol. Mice were deeply anesthetized with ketamine/xylazine (100/10 mg/kg, intraperitoneal) and transcardially perfused with ice-cold Dulbecco’s PBS (DPBS; Corning) followed by freshly prepared 4% PFA. Spinal cords were dissected and post-fixed overnight in 4% PFA at 4°C. For lipid removal, tissues were incubated in 50% CUBIC-L solution (TCI America) at 37°C with gentle shaking and then washed repeatedly in diluted CUBIC-L solution. Samples were subsequently subjected to refractive-index matching using 50% CUBIC-R+(M) solution (TCI America) with gentle rotation until optical transparency was achieved. Refractive-index matching was performed in the dark to preserve fluorescent protein signals.

Cleared spinal cord samples were embedded in 2% agarose and imaged using a Zeiss Lightsheet Z.1 fluorescence microscope equipped with a 5× objective at the UTMB Optical Microscopy Core Facility. Serial optical sections covering the entire spinal cord volume were acquired using identical imaging settings across experimental groups.

### Three-dimensional reconstruction and region segmentation

Raw light-sheet datasets were converted to Imaris-compatible formats using Imaris File Converter (Oxford Instruments) and reconstructed as three-dimensional volumes in Imaris version 10.0.1. For intact mice, spinal cord regions spanning approximately T7–T10 were selected for analysis. In staggered SCI mice, lesion epicenters were identified as regions showing the greatest reduction in neuronal signal intensity. The tissue segment between the two lesion epicenters was defined as the interlesion region. Regions of interest were segmented using Imaris filtering and cropping tools.

### Machine learning-assisted neuronal identification

Automated neuronal segmentation was performed using LABKIT, a machine learning-based image-segmentation plugin implemented in Fiji/ImageJ (https://doi.org/10.3389/fcomp.2022.777728). LABKIT uses sparse manual annotations to train a random-forest pixel classifier for foreground–background segmentation of microscopy images. For each training dataset, approximately 30 representative fluorescent neuronal profiles were manually sampled across different rostrocaudal and dorsoventral locations, imaging planes, signal intensities, and cellular morphologies. Regions corresponding to PRV fluorescence were annotated as foreground, while adjacent non-neuronal structures and background regions with varying fluorescence intensities were annotated as background. These annotations were distributed throughout the image volume to capture spatial and morphological variability within the dataset.

The initial annotations were used to train the LABKIT random-forest classifier. Segmentation results were visually inspected across multiple planes, and additional foreground or background annotations were added at regions of misclassification followed by iterative retraining until neuronal signals could be consistently distinguished from background. LABKIT generates pixel-level feature vectors from image-filter responses and uses these labeled pixels to train its random-forest classifier before applying the classifier to the complete image dataset. The resulting classifier was then applied to the full three-dimensional spinal cord volumes using identical classification settings across experimental groups. Where adjacent neuronal profiles were incompletely separated after classification, individual objects were further refined using region-growing and surface-separation tools in Imaris before three-dimensional quantification.

### Three-dimensional quantification and image analysis

After segmentation refinement, fluorescently labeled neurons within defined spinal cord regions of interest were quantified using the Imaris Surface analysis module. Quantification parameters and threshold settings were held constant across experimental groups. Three-dimensional renderings and neuronal distribution maps were generated in Imaris for analysis and figure preparation.

## QUANTIFICATION AND STATISTICAL ANALYSIS

All quantitative analyses were performed with investigators blinded to experimental group allocation. Statistical analyses and graph generation were performed using GraphPad Prism 10 (GraphPad Software). Data distributions and homogeneity of variance were assessed using the Shapiro–Wilk test for normality and the Brown–Forsythe test for equality of variances, respectively, before selection of statistical tests. For comparisons between two independent groups, unpaired two-tailed *t* tests with Welch’s correction were used for normally distributed data, and Mann– Whitney tests were used for non-normally distributed data. Experiments involving multiple groups or experimental factors were analyzed using one-way or two-way ANOVA, as appropriate. One-way ANOVA was followed by Tukey’s multiple comparisons test, whereas two-way ANOVA was followed by Bonferroni’s or Tukey’s multiple comparisons test, as appropriate. Data are presented as mean ± SD. Statistical significance was defined as *P* < 0.05. Sample sizes and the specific statistical tests and post hoc multiple-comparison procedures used for individual analyses are provided in the corresponding figure legends.

## Acknowledgments

We are grateful to M. Ivannikov from the optical microscopy core at the UTMB for assistance with light sheet imaging. We are also grateful to R. Brantley and W. Russell from the Mass Spectrometry Core at UTMB for assistance with bumetanide concentration detection.

## Author contributions

Q.L. and B.C. conceptualized and designed the study. Q.L. performed the majority of the experiments, analyzed and interpreted the data, and wrote the manuscript. W.L. and J.S. assisted with behavior analysis and PRV tracing. J.S. assisted with immunohistochemistry experiments, SCI surgeries, behavioral assessments, and animal care. A.S. helped design and perform the 3D imaging experiments and contributed to data analysis. T.D. prepared and bred the transgenic mice and assisted with animal care. H.Y.K. assisted with cystometry and optogenetics setup. K.L.V. provided the clinical patient cohort, contributed to clinical data collection and interpretation, and reviewed the manuscript. B.C. supervised the study, interpreted the data, and edited the manuscript. All authors reviewed and approved the final manuscript.

## Supplemental figure legends

**Supplementary Figure 1.**
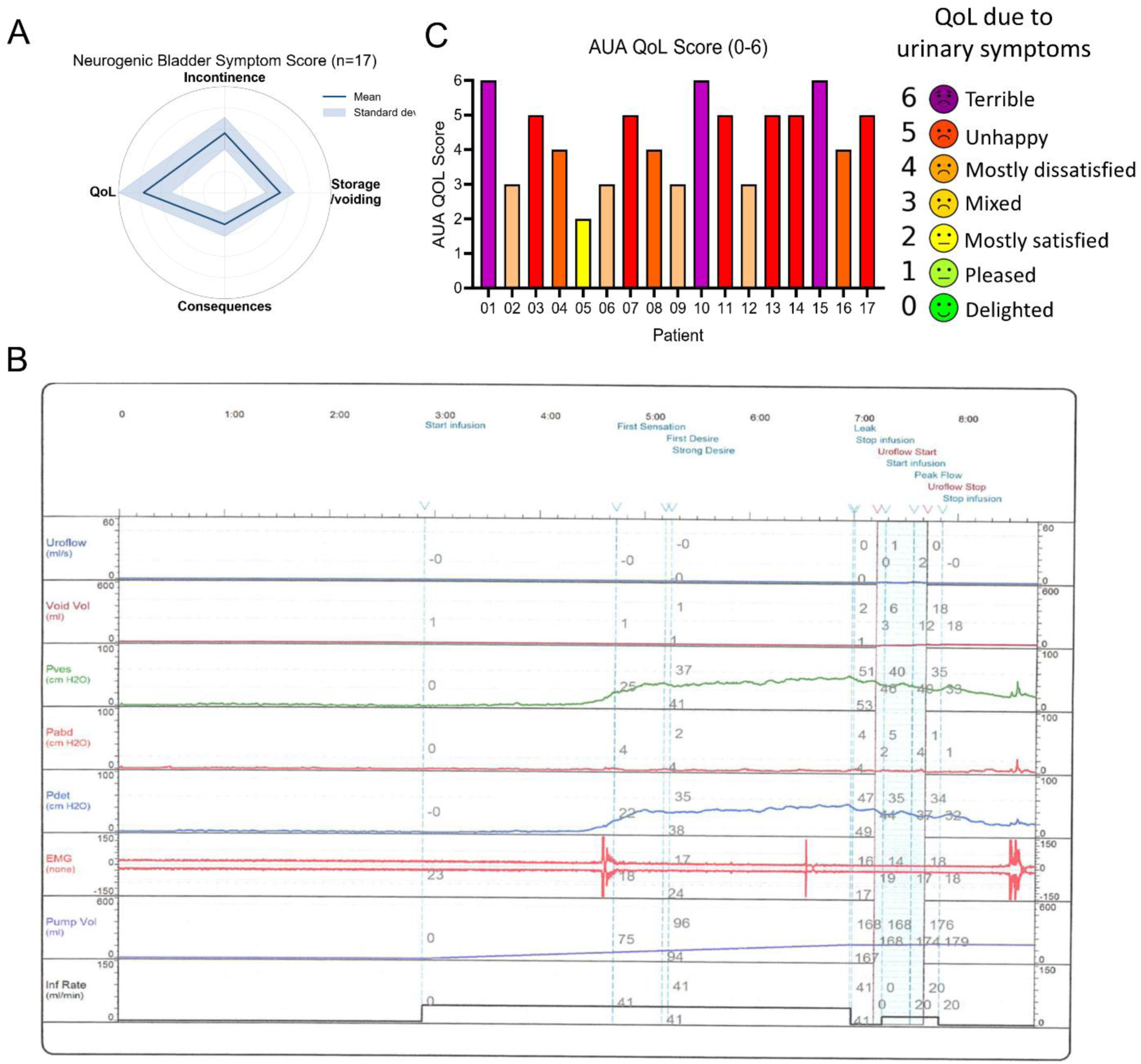
Characterization of neurogenic bladder symptoms in SCI patients. (A) Radar plot of Neurogenic Bladder Symptom Score (NBSS) domain scores, including incontinence, storage/voiding, consequences, and quality of life, in the chronic SCI cohort (n = 17). Data are presented as mean ± SD. (B) Representative multichannel urodynamic recording from a patient with chronic SCI during bladder filling and voiding. (C) Individual American Urological Association (AUA) quality-of-life scores related to urinary symptoms in the SCI cohort, with the corresponding 0–6 response categories indicated.

**Supplementary Figure 2.**
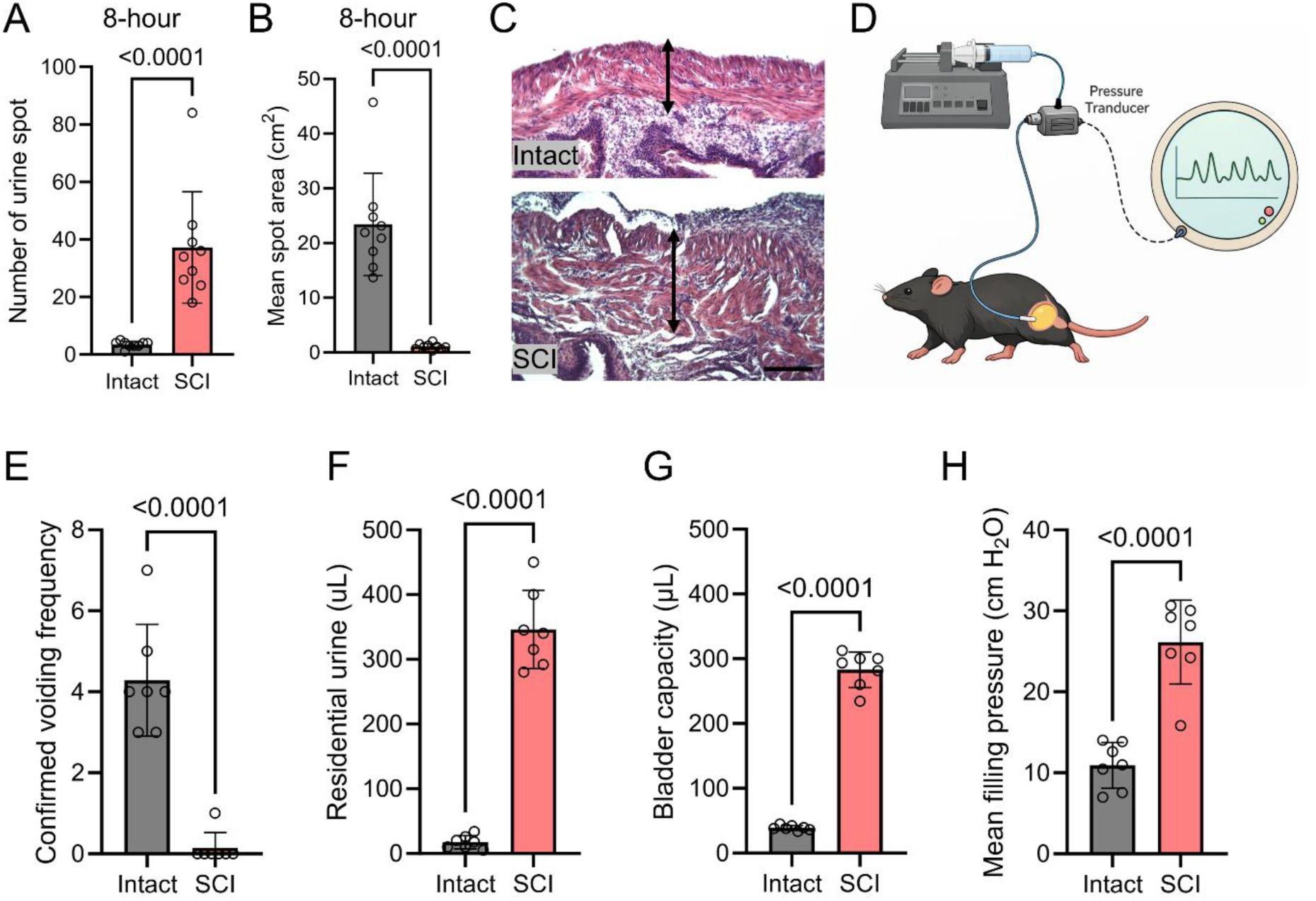
Characterization of neurogenic bladder dysfunction in SCI mice. (A, B) Quantification of urine spot number and mean urine spot area from the 8-h VSOP assay in intact and SCI mice. N = 9, Unpaired t test with Welch’s correction. (C) Representative H&E-stained bladder wall sections from intact and SCI mice. Double-headed arrows indicate bladder wall thickness. (D) Schematic of conscious cystometric recording during bladder saline infusion. (E–H) Quantification of confirmed voiding frequency, residual urine volume, bladder capacity, and mean filling pressure in intact and SCI mice. N = 7, Unpaired t test with Welch’s correction. Data are presented as mean ± SD. Scale bar, 100 μm (C).

**Supplementary Figure 3.**
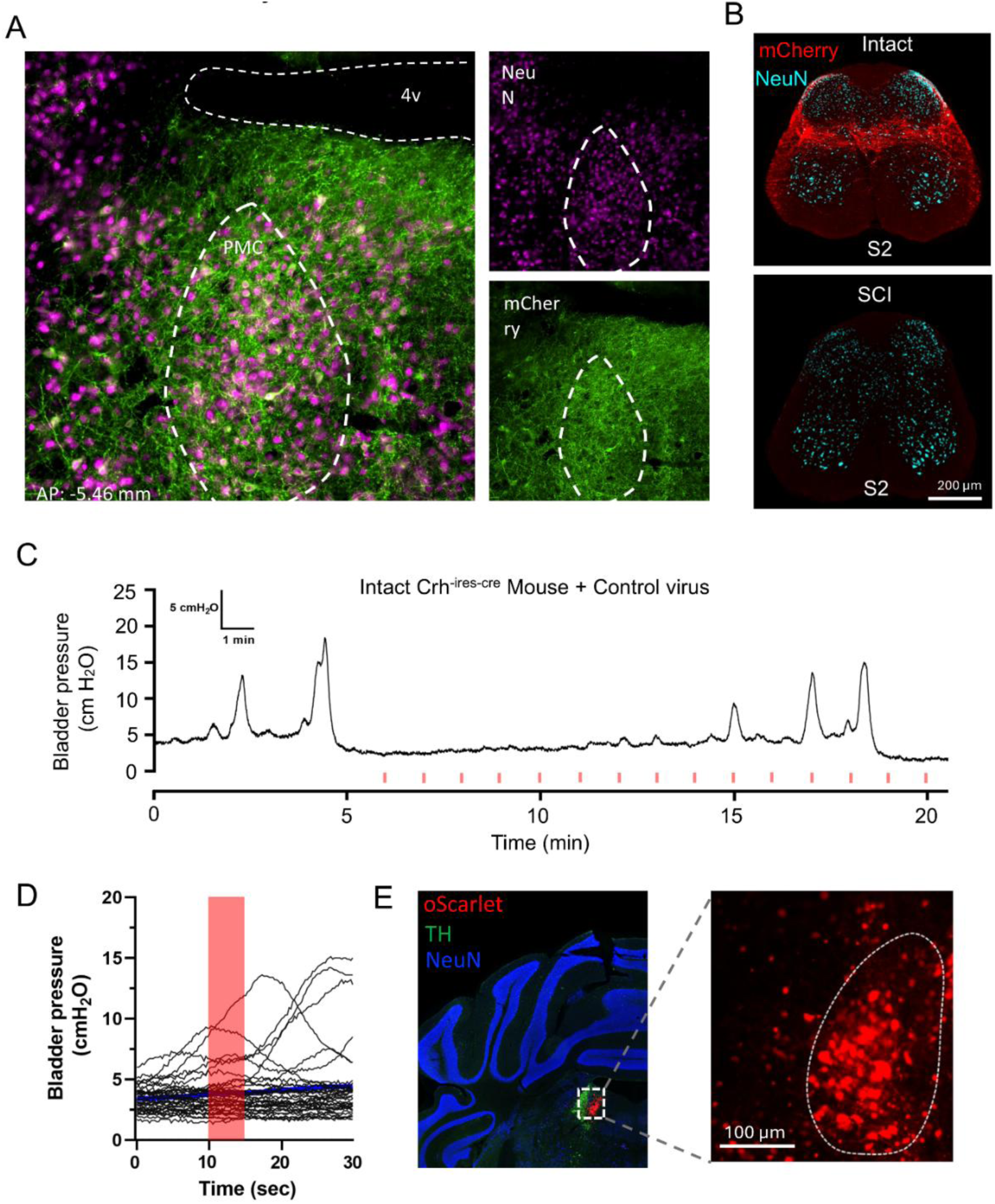
Characterization of PMC descending projections and non-invasive optogenetic controls. (A) Representative image of viral reporter expression in the pontine micturition center (PMC) adjacent to the fourth ventricle (4v). Right panels show NeuN and mCherry channels separately. Dashed outlines indicate the PMC region. (B) Representative S2 spinal cord sections showing PMC-derived mCherry^+^ axons in intact and SCI mice. NeuN is shown in cyan. (C) Representative bladder pressure recording from an intact Crh^IRES-Cre^ mouse injected with control virus during repeated laser stimulation. Red tick marks indicate laser stimulation epochs. (D) Bladder pressure traces aligned to laser onset in control virus–injected mice; the shaded region indicates the laser stimulation period. (E) Representative images of control virus expression in the PMC region. oScarlet is shown in red, TH in green, and NeuN in blue; the dashed outline indicates the PMC. Scale bars are indicated in the panels.

**Supplementary Figure 4.**
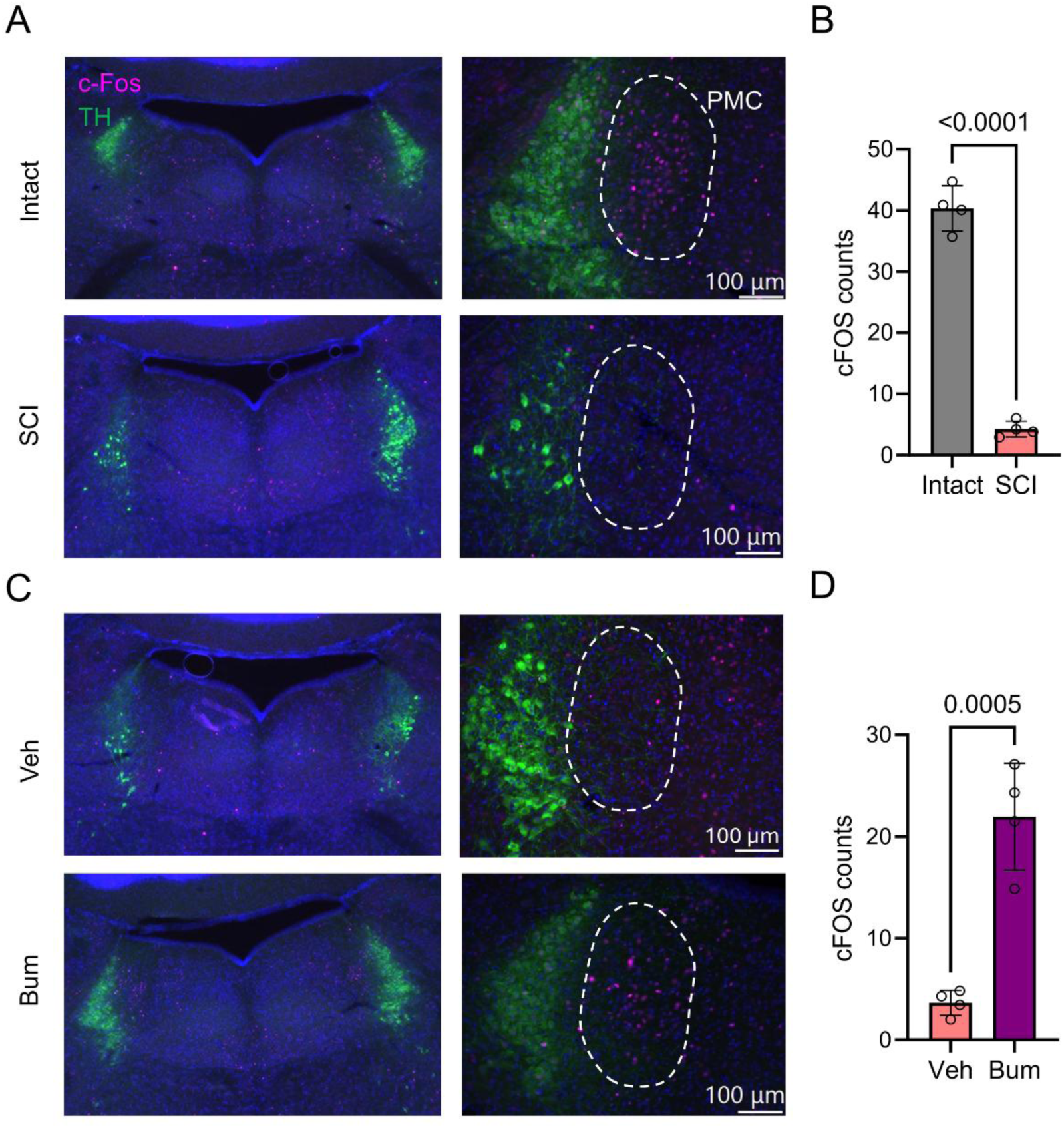
c-Fos induction in the PMC after bladder stimulation. (A, B) Representative c-Fos staining and quantification in the PMC region of intact and SCI mice after bladder stimulation. TH labels the locus coeruleus. White dash indicates PMC area. N = 4, Unpaired t test with Welch’s correction. (C, D) Representative c-Fos staining and quantification in vehicle- and bumetanide-treated SCI mice. N = 4, Unpaired t test with Welch’s correction. Data are shown as mean ± SD. Scale bars, 100 μm.

**Supplementary Figure 5.**
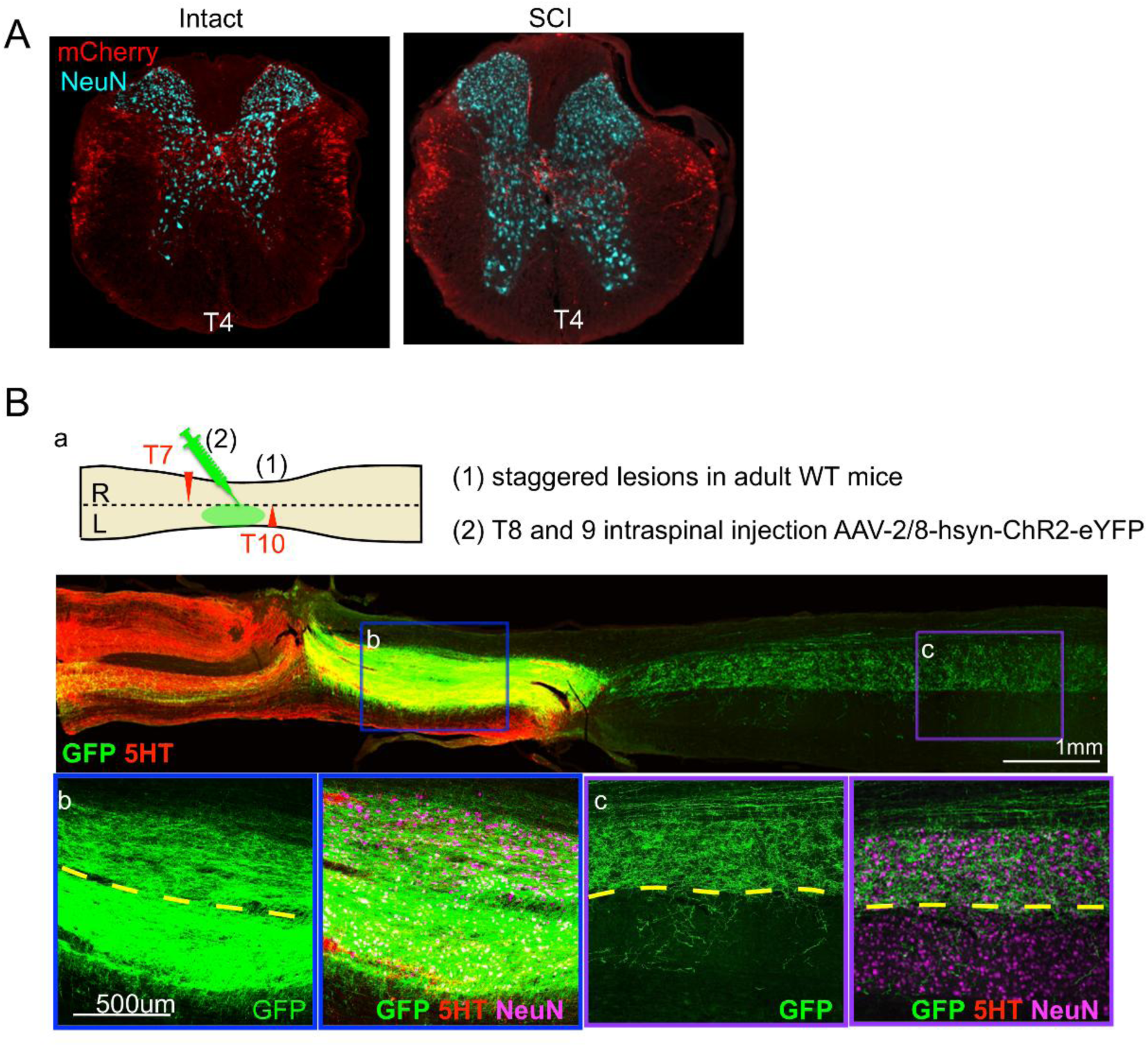
Characterization of spared descending and intraspinal connections in the staggered SCI model. (A) Representative T4 spinal cord sections showing PMC-derived mCherry^+^ descending axons in intact and SCI mice. NeuN is shown in cyan. (B) Schematic of staggered T7 and T10 hemisections followed by AAV2/8-hsyn-ChR2-eYFP injection into the T8–T9 interlesion region. Representative longitudinal spinal cord image shows GFP-labeled intraspinal projections together with 5-HT immunostaining. Boxed regions are shown at higher magnification below with GFP, 5-HT, and NeuN labeling. Dashed lines indicate the spinal cord midline. Scale bars are indicated in the panels.

**Supplementary Figure 6.**
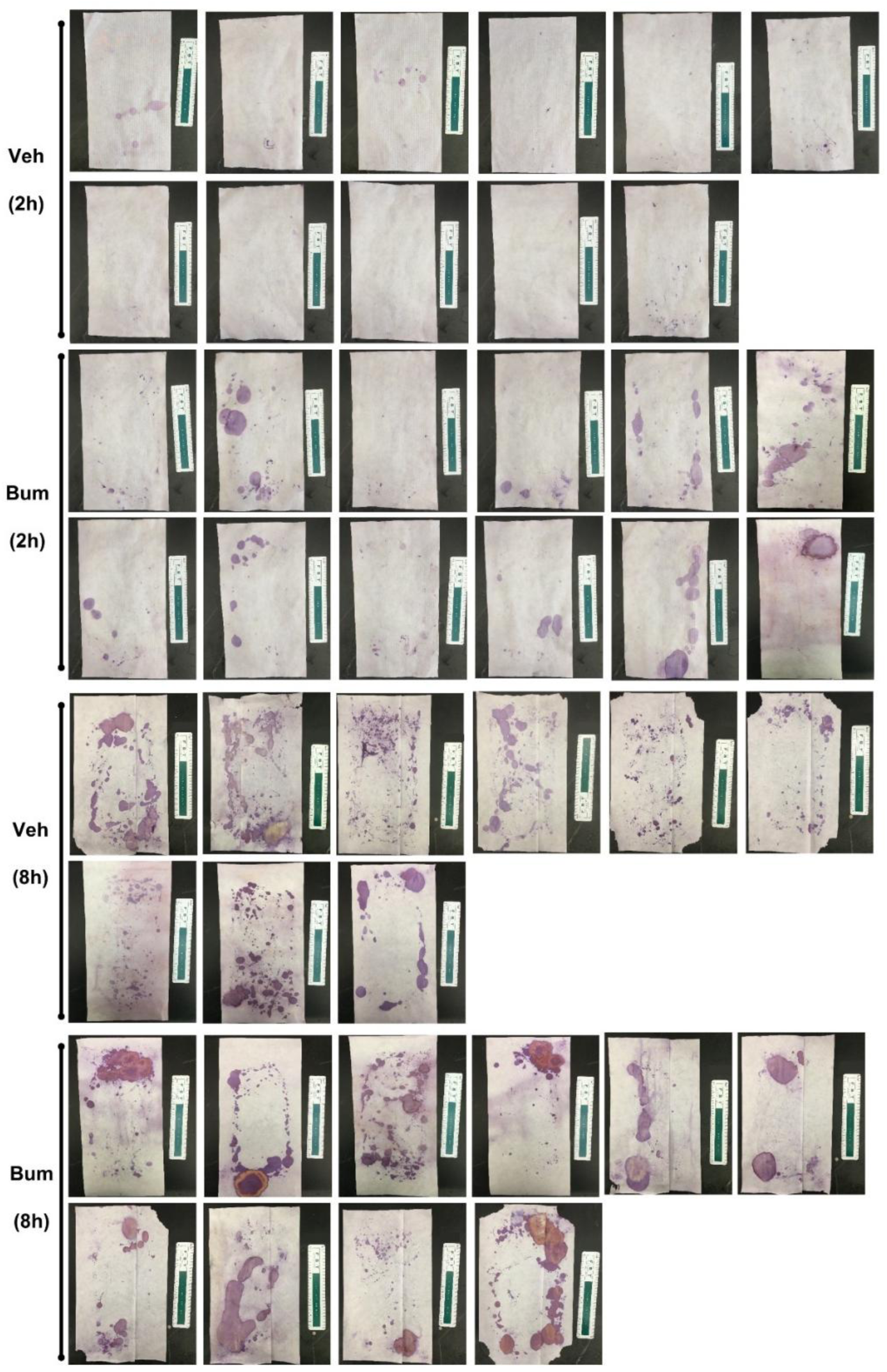
Raw VSOP images from vehicle- and bumetanide-treated SCI mice. Representative raw urine spot on paper (VSOP) images collected from SCI mice treated with vehicle or intrathecal bumetanide during 2-hour and 8-hour recording periods. Individual filter papers from each mouse are shown without image selection or exclusion. Scale bars as indicated on the filter paper ruler.

**Supplementary Figure 7.**
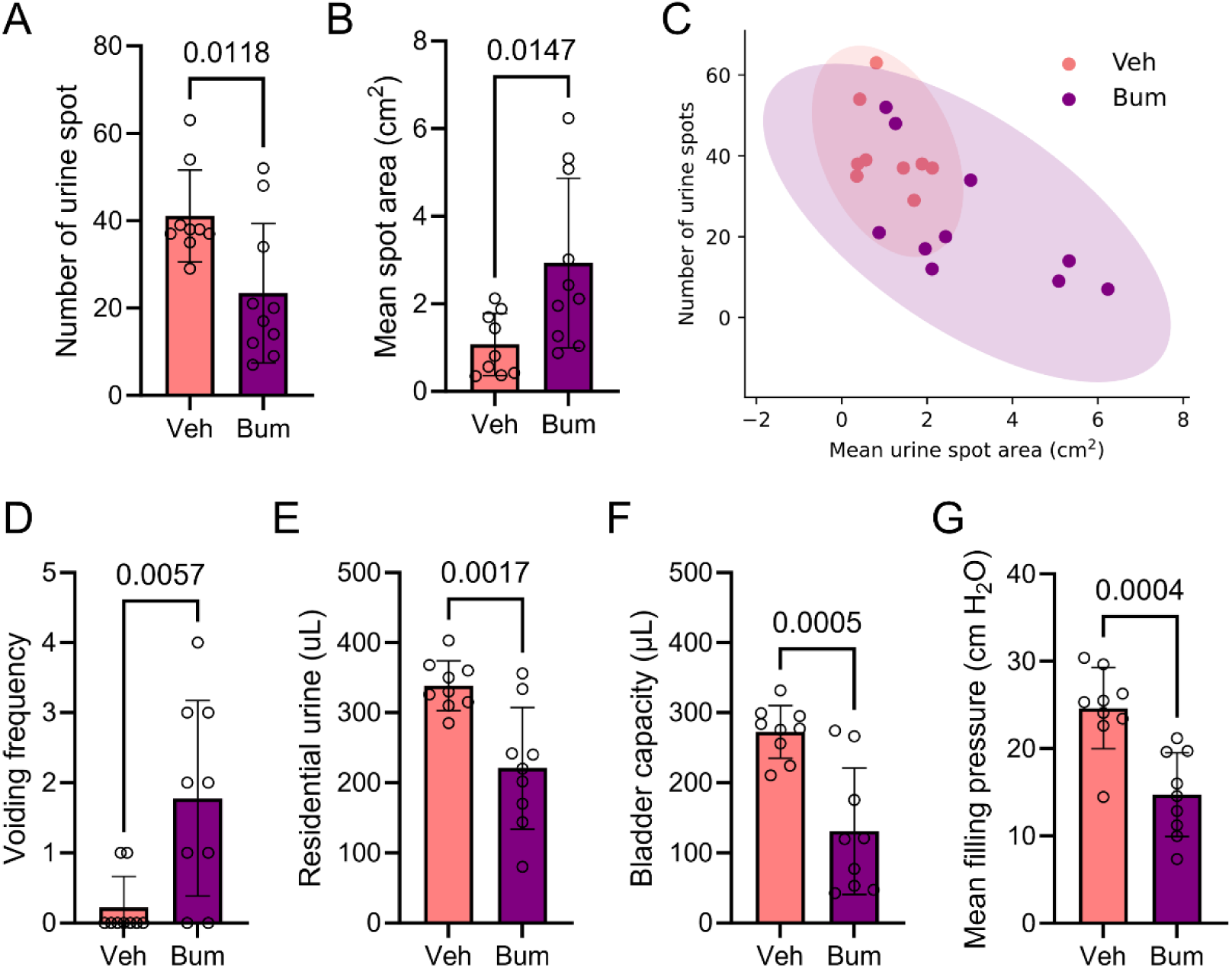
Characterization of intrathecal bumetanide treatment after SCI. (A, B) Quantification of urine spot number and mean urine spot area during the 8-h VSOP assay in vehicle- and bumetanide-treated SCI mice. (C) Bivariate distribution of urine spot number and mean urine spot area for individual vehicle- and bumetanide-treated SCI mice. (D–G) Quantification of confirmed voiding frequency, residual urine volume, bladder capacity, and mean filling pressure in vehicle- and bumetanide-treated SCI mice. Data are presented as mean ± SD. Statistical tests and sample sizes are provided in the corresponding panels and Methods.

**Supplementary Figure 8.**
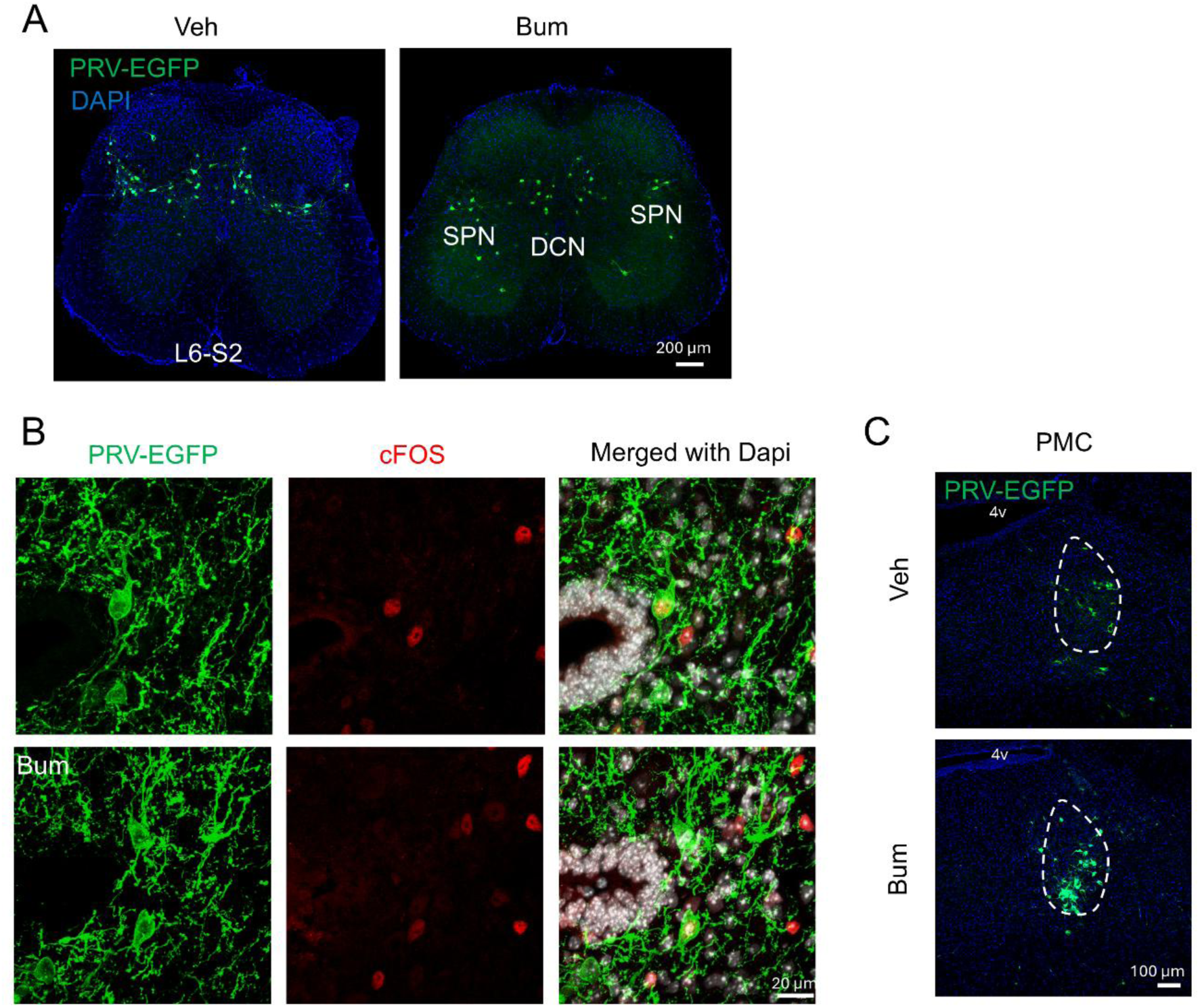
PRV tracing of bladder-connected spinal and supraspinal circuits after bumetanide treatment. (A) Representative L6–S2 spinal cord sections showing PRV-EGFP labeling in vehicle- and bumetanide-treated SCI mice. DAPI labels nuclei; SPN, spinal parasympathetic nucleus; DCN, dorsal commissural nucleus. (B) Representative images of PRV-EGFP and c-Fos labeling in the T8–T9 interlesion region of vehicle- and bumetanide-treated SCI mice. DAPI is shown in the merged images. (C) Representative PRV-EGFP labeling in the pontine micturition center (PMC) of vehicle- and bumetanide-treated SCI mice. Dashed outlines indicate the PMC region; 4v, fourth ventricle. Scale bars, 200 μm (A), 20 μm (B), and 100 μm (C).

